# An HIV-1 Reference Epitranscriptome

**DOI:** 10.1101/2025.01.30.635805

**Authors:** Michael S. Bosmeny, Adrian A. Pater, Li Zhang, Lydia Larkai, Beverly E. Sha, Zidi Lyu, Masad J. Damha, João I. Mamede, Keith T. Gagnon

**Affiliations:** Dept. of Biochemistry, Wake Forest University, School of Medicine, Winston-Salem, North Carolina, USA, 27101; Dept. of Microbial Pathogens and Immunity, Rush University, Chicago, Illinois, USA, 60612; Division of Infectious Diseases, Rush University Medical Center, Chicago, Illinois, USA, 60612; Dept. of Chemistry, McGill University, Montreal, Canada, H3A, 0G3

## Abstract

Post-transcriptional modifications to RNA, which comprise the epitranscriptome, play important roles in RNA metabolism, gene regulation, and human disease, including viral pathogenesis. Modifications to the RNA viral genome and transcripts of human immunodeficiency virus 1 (HIV-1) have been reported and investigated in the context of virus and host biology. However, the diversity of experimental approaches used has made clear correlations across studies, as well as the significance of the HIV-1 epitranscriptome in biology and disease, difficult to assess. Therefore, we established a reference HIV-1 epitranscriptome. We sequenced the model NL4-3 HIV-1 genome from infected primary CD4+ T cells and the Jurkat cell line using the latest nanopore chemistry, optimized RNA preparation methods, and the most current and readily available base-calling algorithms. A highly reproducible sense and a preliminary antisense HIV-1 epitranscriptome were created, where N^6^-methyladenosine (m^6^A), 5-methylcytosine (m^5^C), pseudouridine (psi), inosine, and 2’-*O*-methyl (N_m_) modifications could be identified by rapid multiplexed base-calling. We observed that sequence and neighboring modification contexts induced modification miscalling, which could be corrected with synthetic HIV-1 RNA fragments. We validated m^6^A modification sites with STM2457, a small molecule inhibitor of methyltransferase-like 3 (METTL3). We find that modifications are quite stable under combination antiretroviral therapy (cART) treatment, in primary CD4+ T cells, and in HIV-1 virions. Sequencing samples from people living with HIV (PLWH) revealed conservation of m^6^A modifications. However, analysis of spliced transcript variants suggests transcript-dependent modification levels. Our approach and reference data offer a straightforward benchmark that can be adopted to help advance rigor, reproducibility, and uniformity across HIV-1 epitranscriptomics studies. They also provide a roadmap for the creation of reference epitranscriptomes for many other viruses or pathogens.

## INTRODUCTION

Epitranscriptomics is the study of post-transcriptional RNA chemical modifications and their functional effects on RNA in biology, disease, and beyond (Cerneckis et al., 2024). While epitranscriptomics is an established field of investigation, it is constantly evolving with the characterization of new modifications in unexpected places (Gao et al., 2025; Yan et al., 2025; Zhang et al., 2021), methods to identify them (Cai et al., 2024; Cerneckis et al., 2024; Ron et al., 2025; Roundtree et al., 2017), and pathways they impact (Cerneckis et al., 2024; Jia et al., 2025; Phillips et al., 2024). Nonetheless, experimental and bioinformatic tools like the m^6^A-Atlas (Liang et al., 2024; Tang et al., 2021) or the MODOMICS database (Boccaletto et al., 2022; Dunin-Horkawicz et al., 2006) can provide useful reference points across studies. Viral RNAs, for example, often possess their own unique epitranscriptomes that can either control viral pathogenesis, such as viral gene expression and replication, or modulate host pathways like the innate immune response (Chen et al., 2021; Phillips et al., 2024; Schultz et al., 2025; Verhamme and Favoreel, 2025).

The RNA modifications of human immunodeficiency virus 1 (HIV-1) represent one of the most well-studied viral epitranscriptomes (Phillips et al., 2024; Wang et al., 2022). Most sequencing-based studies have focused on N^6^-methyladenosine (m^6^A) (Baek et al., 2024; Chen et al., 2021; Lichinchi et al., 2016; Mishra et al., 2024; Pereira-Montecinos et al., 2022; Tirumuru et al., 2016; Tsai et al., 2021). Other modifications, including 5-methylcytosine (m^5^C) (Courtney et al., 2019; Huang et al., 2023), N^4^-acetylcytidine (ac^4^C) (Tsai et al., 2020), pseudouridine (psi or Ψ) (Martinez Campos et al., 2021), inosine (Sharmeen et al., 1991) and 2’-*O*-methylation (2’-*O*-Me or N_m_) (Ringeard et al., 2019) have also been investigated but are less understood. It is currently difficult to establish a clear consensus on the RNA modification landscape of HIV-1, especially beyond m^6^A, due to the diversity of experimental methods (Bosmeny et al., 2025). Several groups have mapped suspected regions for a single modification type at a time, often by an antibody-based method, but these do not have single-nucleotide resolution (Kennedy et al., 2016; Lichinchi et al., 2016; Tirumuru et al., 2016). Other techniques that do have single-nucleotide resolution, such as bisulfite sequencing for m^5^C detection (Huang et al., 2023) or m^6^A-SAC-seq for m^6^A (Mishra et al., 2024), can sometimes be challenging to execute and interpret.

Having a standardized methodology for identifying HIV-1 RNA modification that produces consistent results, can be utilized for multiple modifications, is straightforward to implement, and can be correlated across multiple studies as a benchmark would help advance the field (Bosmeny et al., 2025). Here we establish a reference methodology and epitranscriptome for HIV-1 using a common model virus genome, NL4-3, and the Jurkat T cell line. In addition, two RNA library preparation methods, ribosomal RNA (rRNA) depletion and poly-A selection, are combined to provide more consistent coverage followed by nanopore direct RNA sequencing (dRNA-seq) with the latest flow cell chemistry. The most up-to-date base calling algorithms from Oxford Nanopore Technologies (ONT) are applied for multiplex base-calling of m^6^A, m^5^C, psi, inosine, and 2’-*O*-methyl (2’-*O*-Me). We report miscalling errors with current ONT algorithms due to unique sequence and modification contexts and provide baseline subtraction and correction methods to address them. This resulted in a refined modification map of a sense and preliminary, less-explored, antisense epitranscriptome of HIV-1 that can serve as benchmark references. We validated m^6^A modifications using small molecule inhibition of methyltransferase-like 3 (METTL3) and synthetic HIV-1 RNA fragments. We applied our approach to mapping modifications in Jurkat cells treated with physiological levels of current first-line combination antiretroviral therapy (cART) treatment, primary CD4+ T cells, virions from CD4+ T cell culture, and CD4+ T cells from people living with HIV-1 (PLWH), which indicated conservation of most modifications. We also identified differential modification patterns for spliced transcripts versus unspliced HIV-1 RNA using dRNA-seq data. Together, these results provide an initial reference HIV-1 epitranscriptome for multiple modifications and offer a roadmap for creating reference epitranscriptomes using state-of-the-art ONT nanopore sequencing.

## RESULTS

### Direct RNA sequencing provides multiplexed base modification calling on HIV-1 viral RNA

To create an accessible and reproducible methodology for a reference HIV-1 epitranscriptome, we selected a commonly used NL4-3 HIV-1 genome and Jurkat T cells. We also selected the Oxford Nanopore Technologies (ONT) nanopore-based direct RNA sequencing (dRNA-seq) platform. Nanopore sequencing is a cost-effective, commercial technology that is well-supported and can perform direct sequencing of RNA with minimal alteration and without amplification (Jain et al., 2022). Importantly, dRNA-seq has been used previously to sequence HIV-1 viral RNAs (Baek et al., 2024; Honeycutt et al., 2024; Li et al., 2024), albeit with older flow cell technologies and limited baseline correction. However, ongoing improvements in flow cell chemistry, as well as bioinformatic base-calling tools that are continually improving, have made dRNA-seq an increasingly attractive off-the-shelf solution for multiplexed identification of RNA modifications. The combination of these model systems offers an opportunity to establish a benchmark methodology and data set that is robust, reproducible, and accessible to help advance HIV-1 epitranscriptomics studies.

Our initial approach was to infect Jurkat cells with non-replicative HIV-GFPΔEnv/VSV-G virus, which contains a partial replacement of the *nef* gene with GFP that also includes a frameshift in the *env* gene (hereafter referred to as NL4-3-GFP). Total RNA was collected and depleted of rRNA or poly-A selected. The enriched RNA was directly sequenced with an ONT PromethION 2 Solo instrument. We performed base-calling with ONT algorithms to calculate likely modification sites for m^6^A, m^5^C, psi, inosine, and 2’-*O*-Me. Initial experiments were performed in three biological replicates and demonstrated high reproducibility across the NL4-3-GFP HIV-1 genome, in particular for m^6^A (**Figure S1**). To validate modification calling, we treated Jurkat cells with STM2457 at the time of infection. STM2457 is an inhibitor of METTL3 catalysis (Yankova et al., 2021), the primary methyltransferase that writes m^6^A onto RNA. A 30 μM dose of STM2457 was sufficient to reduce detection of all m^6^A modification sites on NL4-3-GFP RNA to less than half their normal value on average (**Figure 1A**). This result supported accurate m^6^A identification by the current ONT algorithm.

**Figure 1.**
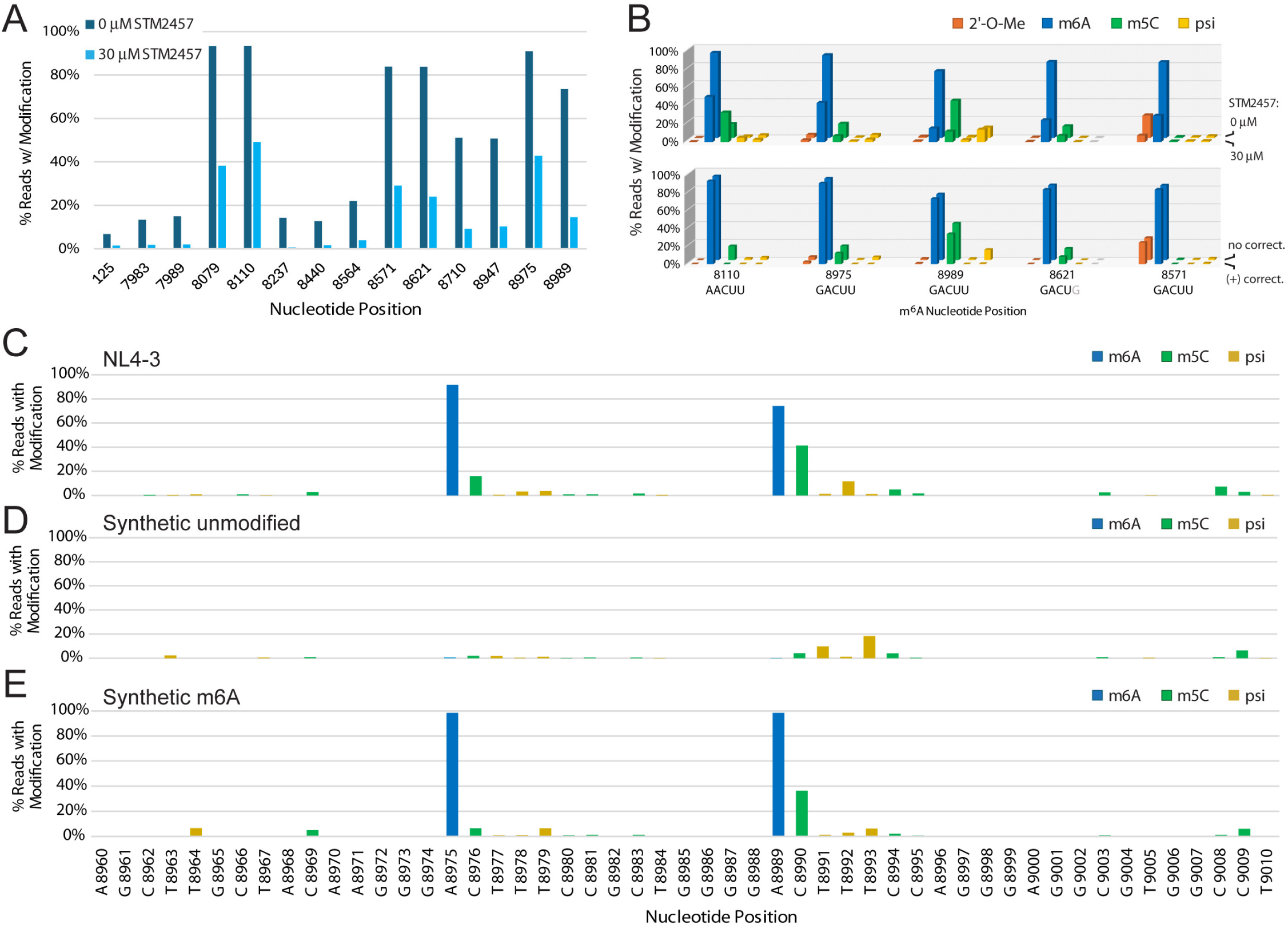
Probing and correction of HIV-1 base modifications called by nanopore direct RNA sequencing. (**A**) Nanopore modification-calling results for HIV-1 viral RNA extracted from Jurkat cell cultures treated with STM2457, a drug that inhibits METTL3 m6A modification activity. The position of each high frequency m^6^A nucleotide in the HIV-1 genome is indicated on the x-axis. (**B**) Comparison of modifications at DRACH motif sites where m^6^A was called at high frequency. The nucleotide position of the m_6_A modification within the DRACH motif is indicated on the x-axis. Upper plot: comparison of Jurkat cell samples infected with HIV-1 without (back row) or with (front row) 30 μM STM2457 treatment. Lower graph: comparison of Jurkat cell sample infected with HIV-1 before (back row) and after (front row) baseline correction. (**C-E**) Comparison of modification calling between NL4-3 from Jurkat cells (**C**) and two synthetic HIV-1 RNA fragments, one unmodified (**D**) and one bearing m^6^A (**E**) at two DRACH motifs. The nucleotide position corresponding to the NL4-3 genome is indicated on the x-axis. All modifications called in panel D are incorrect while modifications called in panel E, besides m^6^A at position 8975 and 8989, are incorrect.

### Background subtraction and correction of HIV-1 RNA modification calls

In our initial data set, an interesting modification pattern was consistently observed at the 3’ end of the HIV-1 transcript, where the majority of m^6^A sites are predicted. METTL3 catalysis of m^6^A strongly prefers a DRACH motif (A/U/G, A/G, A, C, A/U/C). The m^6^A modification itself occurs at the central adenosine, which is always followed by cytosine and then often uridine. Several of these m^6^A modification locations were reported by the base-calling algorithm to also possess significant levels of m^5^C and psi (**Figure 1B**). One possible explanation for such a modification pattern would be miscalling due to a sequence motif context or nearby modifications that influence the base-calling of others. Nanopore flow cells read nucleic acids in five-nucleotide ‘frames,’ or k-mers, which shift one base at a time to generate current signals. This can lead to modification calling artifacts by algorithms during computational interpretation of current signals, especially for uncommon sequence motifs or nearby unaccounted modifications. Unexpectedly, treatment with STM2457 not only reduced the m^6^A calling at these DRACH motifs, but also reduced the m^5^C call counts, suggesting a dependence on m^6^A presence (**Figure 1B**). Conversely, the low levels of psi calls in DRACH motifs did not respond to STM2457 treatment while a 2’-*O*-Me call did at one site, position 8570. Thus, we questioned the legitimacy of m^5^C and potentially other modifications in and immediately adjacent to DRACH motifs, as well as the overall fidelity of multiplexed modification base-calling. To better understand the dependence of m^5^C calls on adjacent m^6^A in DRACH motifs, we chemically synthesized RNA oligonucleotides bearing two adjacent DRACH motifs found in the NL4-3 genome where m^6^A is called, at positions 8975 and 8989. One oligonucleotide contained no modifications while the other contained 100% m^6^A at both positions. When these oligonucleotides were sequenced, we observed an increased frequency of m^5^C miscalling in the DRACH motifs when m^6^A was present (**Figure 1C-E**). Interestingly, psi and 2’-*O*-Me were observed at low but detectable levels in unmodified RNA at certain nucleotide positions, as was m^5^C to a lesser extent, indicating background calling due to sequence context (**Figure 1D**, **Figure S2**).

To address the potential influence of sequence motifs on modification calling at a broader scale, we prepared overlapping multi-kilobase DNA templates from an NL4-3 plasmid (replicative variant without GFP) for *in vitro* T7 RNA polymerase transcription. Fragments were *in vitro* transcribed with standard nucleotide triphosphates and sequenced, revealing very minimal background miscalling for m^6^A, low miscalling for psi, inosine, and 2’-*O*-Me, and more substantial sequence-dependent miscalling for m^5^C. (**Figure S3**). Using the modification calling frequencies from *in vitro* transcribed NL4-3 fragments, we performed a baseline correction (Baek et al., 2024). After correction, the original calling of m^6^A remained nearly unchanged, indicating that the base-calling algorithm is quite accurate (**Figure 2A, Figure S1, Table S1**). In contrast, only a handful of high-confidence calls for m^5^C, inosine, and 2’-*O*-Me modifications were observed across the NL4-3-GFP viral RNAs after correction using a 10% cutoff threshold (**Figures 2B and 2D-E, Tables S2, S4, and S5**). Baseline correction resulted in no pseudouridine calling frequencies above 10%, removing all potentially significant pseudouridine modifications (**Figure 2C, Table S3**).

**Figure 2.**
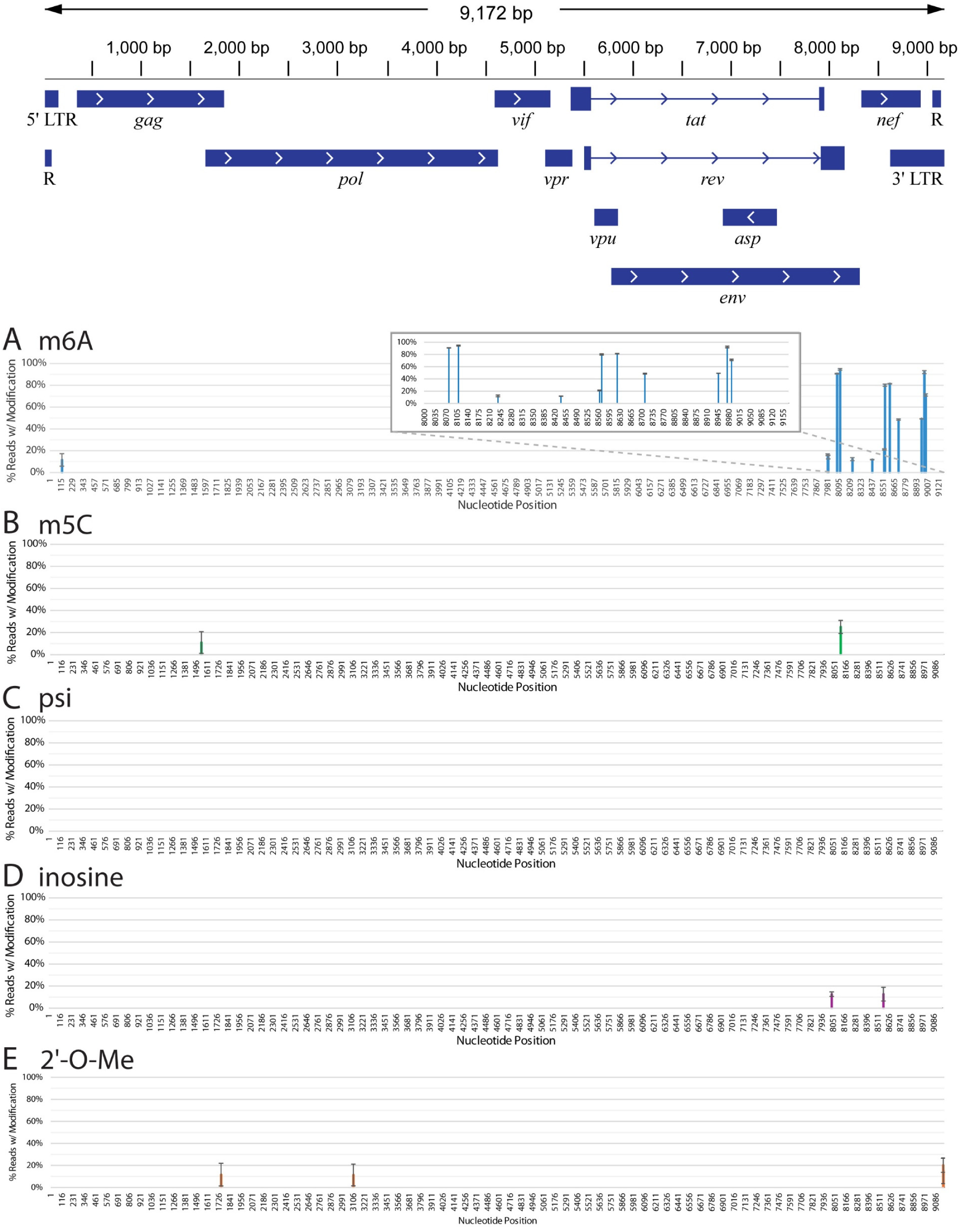
Baseline-corrected nanopore modification calling results for HIV-1 viral RNA from Jurkat cells. The HIV-1 genome architecture is illustrated above. Modifications are shown if, after baseline correction, the average modification frequency was at least 10%. m^6^A (**A**, blue), m^5^C (**B**, green), pseudouridine (psi) (**C**, yellow), inosine (**D**, purple), and 2’-*O*-methylation (**E**, orange). Inset in panel A is a close-up of the 3’ end of the NL4-3 HIV-1 genome where m^6^A is most densely called. Results are the average of three separate biological replicates. Error bars are standard deviation.

To consider the effect of the presence of one modification on the calling of another, the same overlapping multi-kilobase DNA templates were *in vitro* transcribed using modified bases to produce NL4-3 transcripts in which all adenosines had been replaced with m^6^A, all cytosines had been replaced with m^5^C, or all uridines had been replaced with pseudouridine. These heavily modified transcripts were also sequenced and modification frequencies for the other modifications were called. To these new sets of modifications, baseline correction was also applied, to remove false modification calls that would be present without adjacent modifications. For example, the baseline of m^5^C calls from unmodified *in vitro* transcribed RNA was subtracted from the m^5^C modification calls from *in vitro* transcribed RNA with only m^6^A modifications to derive a list of incorrect m^5^C modification calls resulting from nearby true m^6^A modifications. Several trends in the current generation of Nanopore modification calling algorithms are exemplified in these analyses (**Figure S4**). m^6^A, psi, and inosine can be falsely called sporadically by other nearby modifications, but there is not a strong pattern in these calls. On the other hand, m^5^C is frequently miscalled immediately 3’ of m^6^A in DRACH motifs. Calling of 2’-*O*-Me appears to be the lowest accuracy when other modifications are overlapping or adjacent. For example, when m^6^A is present, A_m_ on the same nucleotide or a G_m_ immediately 5’ of the A in the DRACH motif will frequently be miscalled. When m^5^C is present, G_m_ or C_m_ immediately 5’ can be incorrectly called. When psi is present, an A_m_ or C_m_ directly 5’ can also be miscalled. Using the current ONT algorithm, 2’-*O*-Me calls directly 5’ of another modification should be considered with caution. A prominent example is a G_m_ call at position 8570, directly 5’ of a called m^6^A. When STM2457 is used to reduce the amount of m^6^A present, G_m_-8570 also decreases substantially. Thus, this was considered a false call and removed from the final list of corrected modifications. Similarly, m^5^C modification calls at 8976 and 8990 are both 3’ of m^6^A modifications and have been removed. Well-defined sequences, especially viral transcriptomes and RNA virus genomes, should be amenable to these methods of baseline correction.

### Features of the corrected HIV-1 epitranscriptome

Several observations can be noted for the different modifications identified in NL4-3-GFP viral RNAs. For m^6^A, nearly all high-frequency modifications are found on the 3’ end of the genome, in approximately the last thousand bases (**Figure 2A, Table S1**). These include a pair at positions 8079 and 8110, which occur in the region where *rev* and *env* genes overlap, and a trio at 8947, 8975, and 8989 in the 3’ LTR just beyond the end of the *nef* gene. Evidence for m^6^A modifications in these areas is well-documented, with previous MeRip-Seq and PA-m^6^A-Seq experiments showing high probabilities of m^6^A in these regions (Courtney et al., 2019; Kennedy et al., 2017; Lichinchi et al., 2016; Pereira-Montecinos et al., 2022; Tirumuru et al., 2016). One group used silent mutations to remove A nucleotides, either individually or altogether, at positions A8079G, A8975C, and A8989T (Baek et al., 2024). The triple mutation resulted in reduced HIV-1 protein expression. There is also a trio of m^6^A modifications in the middle of the *nef* gene (8571, 8621, and 8710), but it is possible that these methylations are artificially induced. The NL4-3-GFP genome possesses a GFP tag that replaces approximately the first one hundred bases of the *nef* gene and the frequency of these modifications are dramatically reduced when a non-GFP genome is used, which we describe later. Finally, there is a single m6A modification (A125) that is present in the 5’ LTR. While it does not exhibit a high level of modification, it is notable for being the only reproducible and significant m^6^A not on the 3’ end of the genome. A previous study suggested an m^6^A in this region but lacked single-nucleotide resolution (Tirumuru et al., 2016).

After baseline correction, only two m^5^C modifications emerged (**Figure 2B**). Position 8114 in the *rev*/*env* region is the more consistent of the two modification calls. It is approximately 15-25% in all samples (**Table S2**). Position 1551 in the *gag* gene, by contrast, is more varied in m^5^C calling frequency from sample to sample. Variability between samples suggests the possibility that this site is more dynamically regulated or sensitive to cellular signals.

The ONT base-calling algorithm identified a large number of potential psi modifications when uncorrected (**Figure S1**). However, most of these were also called in the unmodified *in vitro* transcribed viral genome fragments (**Figure S3**), indicating they are primarily incorrect calls. After removing background, there were no pseudouridine calls that met our 10% threshold cutoff (**Figure 2C**). A lower threshold of 5% was used to generate a list of potential psi modification sites (**Table S3**) for further investigation. Like m^5^C, there is higher variance in psi calling frequency from sample to sample.

Even prior to background subtraction, inosine modifications were called infrequently (**Figure 2D**). Most sites ranged between 10-20% modification. After background subtraction, only two positions reached a 10% threshold. Inosine may only be present on the HIV-1 sense transcript in low abundance and may not be a significant target of adenosine deaminases under these conditions. The highest frequency inosine modifications were 8037 (*env*) and 8568 (*nef*). Interestingly, both of these modifications decreased in the STM2457-treated sample, despite not being immediately adjacent to an m^6^A (**Table S4**). The inosine called at position 8568 is closely located between an m^6^A at position 8564 and 8571. It is possible that the k-mer frame of the sequencer produces an incorrect call despite being a few bases away from m^6^A on either side. In contrast, the nearest m^6^As to position 8037 are quite distant, at 7989 and 8079. Thus, more investigation should be performed to understand the potential co-dependence of inosine on STM2457 treatment.

Calling of 2’-*O*-methylation is the latest algorithm to be added to Nanopore dRNA-seq. At present, it seems to be less reliable than the other algorithms. Nonetheless, there are four sites where 2’-*O*-Me calling passes a 10% threshold: 1749 (*gag*/*pol*), 3107 (*pol*), 9162 (3’ LTR), and 9164 (3’ LTR) (**Figure 2E, Table S5**). At each site, the frequency of methylation appears variable. However, we observed that samples from Jurkat cells grown at the same time have similar frequencies of methylation, suggesting sensitivity to cellular conditions.

### Transcript splice isoforms possess distinct RNA modification frequencies

To examine potential differences in modification patterns between spliced HIV-1 transcripts, a sequencing run of HIV-1 from Jurkat cells with the highest number of full-length reads was used to sort reads into spliced transcript isoform bins using the FLAIR software package (Tang et al., 2020). HIV-1 transcripts can be full-length, or large sections spliced such as the *gag*/*pol* region (Emery and Swanstrom, 2021). There are two splice site exons, S1 and S2, which can optionally be included in splice variants (Emery and Swanstrom, 2021). Not including new combinations produced by adding one or both splice donors, there are approximately six primary isoforms aside from the full-length, unspliced form (Emery and Swanstrom, 2021; Sertznig et al., 2018) (**Figure 3**).

**Figure 3.**
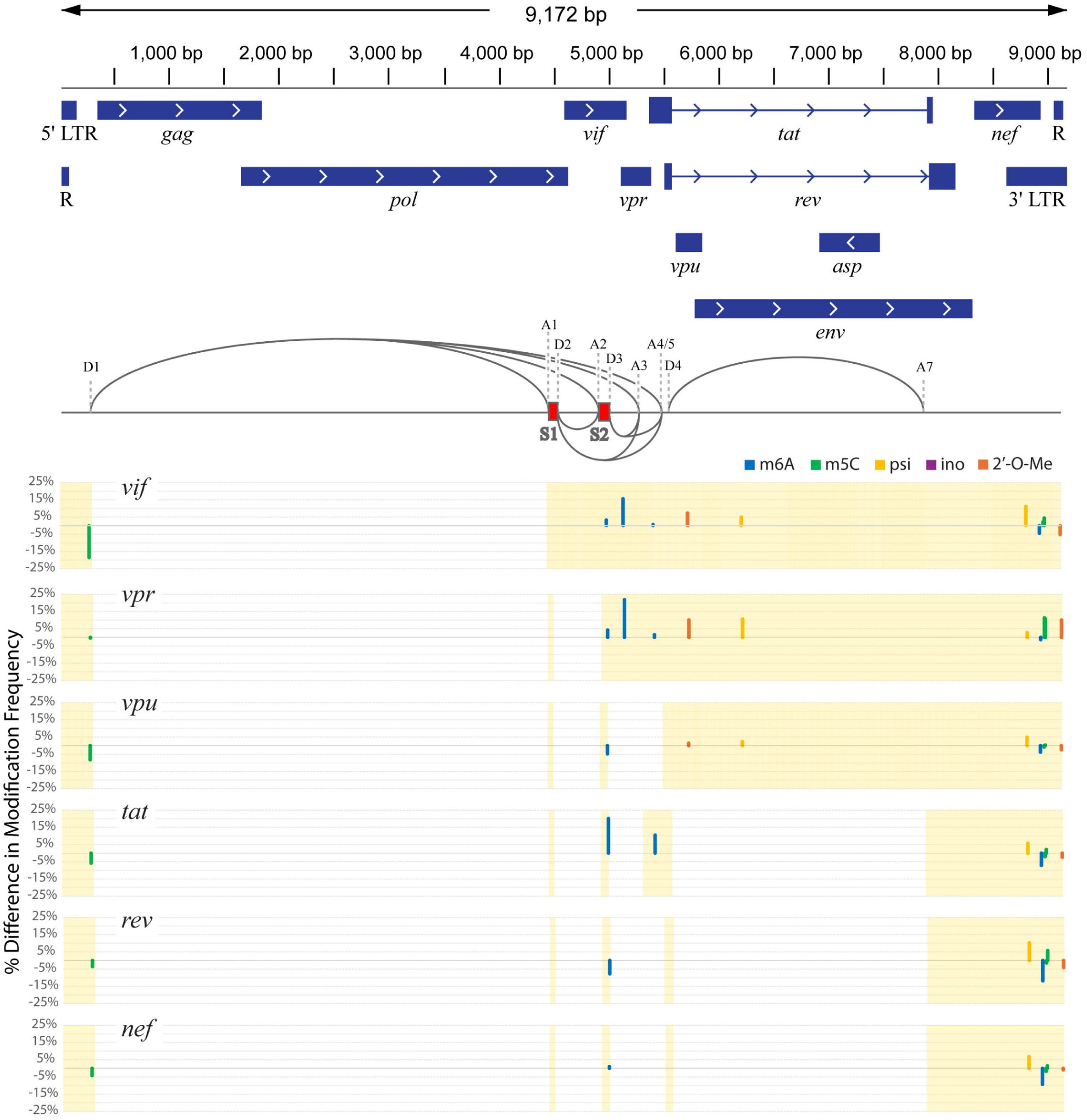
Differential modification frequencies in HIV-1 RNA splice isoforms. Comparison of modification frequency in each splice isoform as compared to the full-length (unspliced) isoform. X-axis represents individual nucleotides in the HIV-1 genome. Y-axis represents percentage of reads that contained the mutation of interest for that nucleotide, relative to the unspliced form. Bar color indicates modification type: m^6^A (blue), m^5^C (green), pseudouridine (yellow), inosine (purple), and 2’-*O*-methyl (orange). Only modifications with a difference of 10% from the unspliced RNA in at least one isoform are shown. Shaded background shows portion of the full transcript retained in the spliced isoform. Splice isoforms are subsets of direct-RNA sequencing sample 7C. The A6 splice site is exclusive to the HXB2 genome and therefore not indicated here.

After sorting, reads were further split and grouped into each splicing variant and modifications were called independently for each variant, including background subtraction as described earlier. The full-length, unspliced read, which codes for the whole genome, including *gag*/*pol*, was used as a comparative baseline. Modifications in which there was at least a 10% difference in modification frequency at that nucleotide between the baseline and one of the other splice isoforms are plotted in **Figure 3** and listed in **Table S6**. Several trends in differential spliced transcript modifications are notable. First, many are near the splice sites. For example, both *vif* and *vpr* splice isoforms have an m^6^A modification at position 5160, a 2’-*O*-Me at 5750, and a psi at 6242 that are increased as compared to the unspliced transcript. Similarly, another m^6^A modification at position 5434 is near the beginning of *tat* and increased in *tat* splice isoforms compared to the unspliced transcript. Interestingly, there is an m^5^C modification near the 5’ LTR at nucleotide position 265 that is of higher frequency in the unspliced transcript than any spliced isoform. One differential modification, m^6^A at 5006, is in the S2 splice donor, which is be incorporated into multiple splice isoforms and most abundant in the *tat* spliced transcript isoform.

The remainder of differential modification calls are in transcript 3’ LTRs. These include m^5^C at positions 9008 and 9019, both of which are called most frequently in the *vpr* splice isoform. A psi at position 8851 is nearly absent in the unspliced form but most frequently called in *vif* and *rev* isoforms. A 2’-*O*-Me at position 9166 (adjacent to constitutively methylated bases at positions 9162 and 9164) is most frequently called in the *vpr* isoform and least in the *vif* isoform. Most of these modifications are of relatively lower frequency, such that they did not reach the 10% average baseline-corrected threshold. However, the m^6^A at position 8975 is very prominent. All splice isoforms combined yielded a calling frequency of 89%, the unspliced form had the highest at 94% and the *rev* isoform had the lowest at 82%.

To explore whether any of these modifications might have an effect on splicing, the list of modifications was compared against a list of known splicing enhancers or silencers (Helene et al., 2018). Those modifications that fall within a splicing regulatory element are listed in **Table S7**. The m^5^C modification called at position 265 is within ESE U5, which is speculated to be bound by SR protein SC35 (SRSF2) to enhance the activation of splice donor D1 (Asang et al., 2012). m^5^C is most frequently expressed at this position in the unspliced isoform, suggesting possible influence on SRSF2 binding and inhibition of splicing. The m^6^A modification at position 5006 is within HIVE3D3, a motif shown to be bound by high mobility group A protein 1a (HMGA1a) (Tsuruno et al., 2011). This binding leads to an increase in the amount of *vpr* splice isoform made, as it promotes the activity of splice acceptor A2 and silences the activity of D3. However, for our sample, there does not seem to be any impact of this m^6^A modification on this D3 splice activity since the *vpr* splice isoform has similar m^6^A levels at 5006 as other splice isoforms. Finally, the m^6^As called at positions 7983 and 7989 are not significantly differentially expressed between splice isoforms despite falling within the ESE-3 splice enhancer. ESE-3 enhances the splicing of splice acceptor A7, which is utilized by the *tat*, *rev*, and *nef* splice isoforms (Staffa and Cochrane, 1995). Thus, these modifications are unlikely to play a role in the regulation of their cognate splice site.

### A novel HIV-1 antisense epitranscriptome

Nanopore dRNA-seq and analysis also enables differentiation of transcript orientation, which allows for modification calling on antisense transcripts like those utilized by the *asp* gene. However, antisense transcripts are very low in abundance, especially outside of the *asp* gene. To interrogate the antisense genome, we bioinformatically combined antisense reads from eleven Jurkat cell samples. This produced coverage ranging from 10X to 50X across the entire HIV-1 antisense transcriptome. Because these reads have lower coverage and are combined from multiple samples, results may be more variable and should be considered more preliminary. To reduce the background noise, we performed correction with unmodified *in vitro* transcribed antisense RNA fragments and only considered modification calling when at least 10x coverage was present with a calling frequency of >20% (**Table S8**). However, modification calling within the *asp* gene (positions 6910 – 7479) was set to a lower cutoff threshold of 10% frequency due to higher average coverage, its gene status, and previous investigation of this *asp* transcript modification (Estevez et al., 2022). Interestingly, there were no m^6^A calls >20% frequency in the antisense epitranscriptome and only one at just under 20% in the *asp* gene (**Figure 3A**). Conversely, inosine modifications, which were rare on the sense epitranscriptome, occur at multiple positions in the antisense direction with modification frequencies at 20-30%. The modification called with highest frequency was inosine at positions 95 and 118, with 28% and 30%, respectively (**Figure 3A and Table S8**). Within the *asp* gene itself, the most prominent modification is inosine at 6911. The modification called with highest frequency across the entire antisense transcriptome was psi at 1093. All five modifications that can be currently called with ONT algorithms, m^6^A, m^5^C, psi, inosine, and 2’-*O*-Me, were previously identified by mass spectrometry at low or moderate abundance in *asp* gene antisense transcripts (Estevez et al., 2022). Modifications to the antisense transcripts have not been previously site-specifically mapped and may play a role in the processing or function of the HIV-1 antisense transcriptome.

### RNA preparation and cART treatments do not alter modification calling

During sample preparation, two methods of RNA enrichment were performed, a bead-based poly-A selection method and an RNase H-based ribosomal RNA (rRNA) degradation method. Each method yielded a different configuration of RNA fragments for sequencing, allowing a greater diversity of reads (**Figure S5A**). Poly-A selection produced relatively long reads from dRNA-seq that usually began at the 3’ end of the viral genome, resulting in high 3’ end coverage but low coverage across the middle and 5’ regions. On the other hand, rRNA depletion resulted in smaller RNA fragments that were well-distributed across the NL4-3-GFP genome. In addition, we observed that poly-A selection alone does not capture host non-coding RNAs efficiently while rRNA depletion does not cover protein coding mRNAs sufficiently (**Figure S5B**). Thus, by combining both methods, either experimentally before sequencing or bioinformatically after, coverage across viral and host transcriptomes was optimized.

There is the possibility that these selection methods could result in modification calling bias due to RNA species enrichment. Therefore, we compared modified base calling between these two methods. The same sample was prepared by both methods separately and sequenced independently. The 3’ end of the HIV-1 genome was compared for modifications, as this region received reasonable coverage under both methods. Our analysis shows little difference between methods when calling modifications for HIV-1 (**Figure S5C**). Therefore, it is anticipated that using one or both RNA preparation methods should not significantly alter modification calling for HIV-1.

We also sought to examine the effect of combination antiretroviral therapy (cART). Three replicates of Jurkat cells, infected with NL4-3-GFP virions, were treated with cART prior to RNA collection. These samples were processed and sequenced as before. We observed no obvious differences in modification calling between samples with and without cART treatment (**Figure 4B**). This suggests that cART does not substantially perturb the enzymatic modification pathways in these cells and is not a clear modifier of the HIV-1 epitranscriptome.

**Figure 4.**
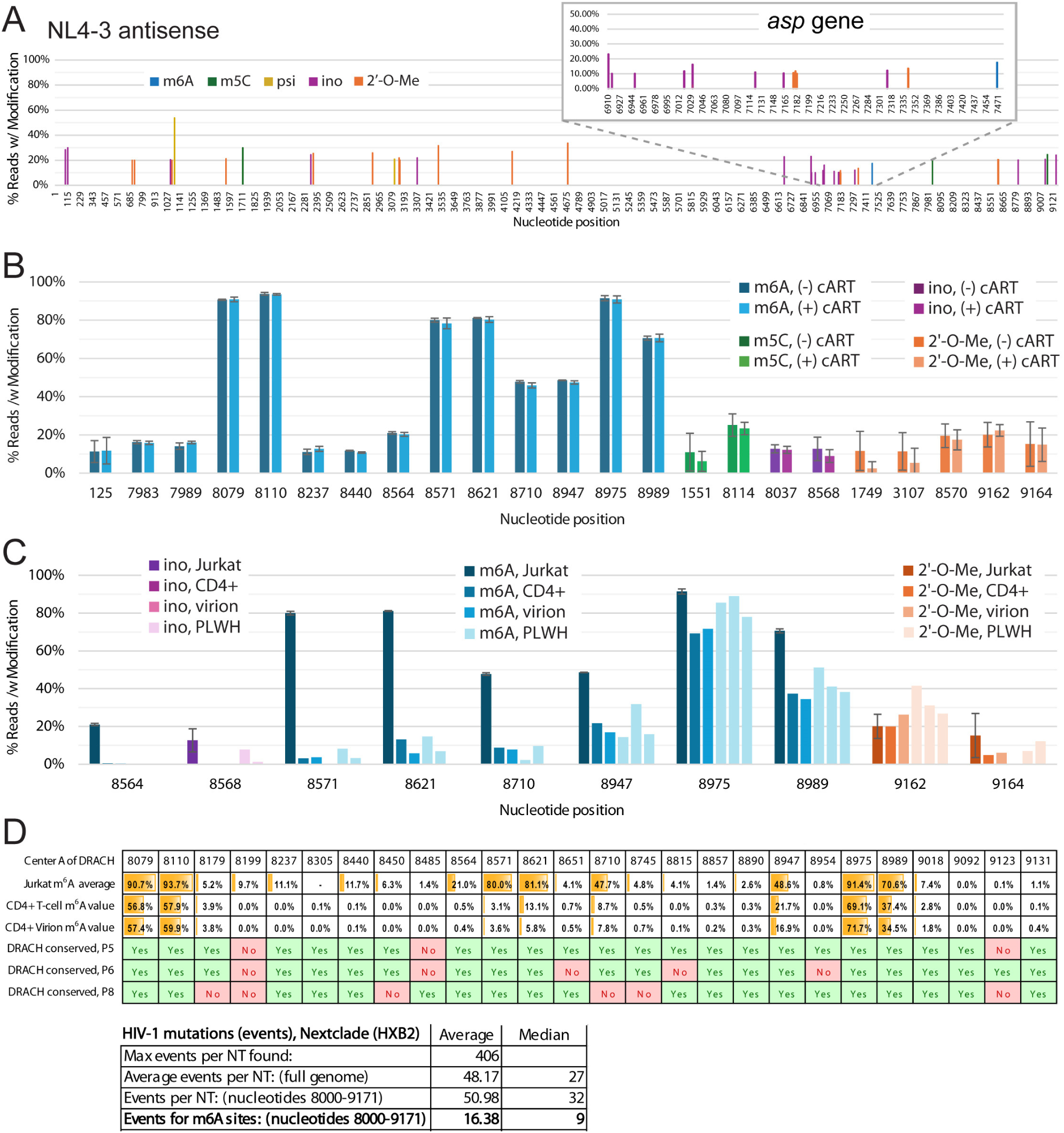
A preliminary HIV-1 antisense epitranscriptome, the effect of cART and cell type on HIV-1 modification calling, and conservation of m^6^A in HIV-1 genomes from PLWH. (**A**) Nanopore modification-calling for HIV-1 antisense RNA from Jurkat cells. m^6^A (blue), m^5^C (green), pseudouridine (yellow), inosine (purple), and 2’-*O*-methylation (orange). Inset shows a close-up of the *asp* gene. (**B**) Nanopore modification-calling results for HIV-1 RNA taken from Jurkat cells treated without (darker color) or with (lighter color) cART treatment. Error bars represent the standard deviation from three separate biological replicates. (**C**) Nanopore modification-calling results for HIV-1 RNA taken from Jurkat cells, primary CD4+ T cells infected *in vitro,* supernatant of primary CD4+ T cells infected *in vitro*, and CD4+ T cells from PLWH samples. In each nucleotide position cluster, the first bar is Jurkat cell samples infected with HIV-1, the second bar is CD4+ T cells from healthy donors and infected with HIV-1, the third bar is the supernatant from the CD4+ T cells from healthy donors, and the fourth, fifth, and sixth bars are samples taken from CD4+ T cells from PLWH donors. (**D**) Top: comparison of m^6^A modifications called from HIV-1 RNA from Jurkat cells against preservation of the DRACH motifs for these m^6^A modifications sequenced from three PLWH samples. Bottom: analysis of conservation of known m^6^A modification sites in a larger dataset of HIV-1 mutations. Positions containing m^6^A between nucleotide positions 8000 and 9171 were identified and the average and median events for these positions are reported.

### The HIV-1 epitranscriptome in CD4+ T cells from people living with HIV

To assess the *ex vivo* and *in vivo* epitranscriptome of HIV-1, including derived from people living with HIV (PLWH), we first performed sequencing of HIV-1 viral RNA from cultured primary infected CD4+ T cells, which were isolated from whole-blood of healthy donors. CD4+ T cells were activated and subsequently infected with replicative wild-type NL4-3 HIV-1. The supernatants of these cultured CD4+ T cells were also collected for HIV-1 virions. We then harvested CD4+ T cells from whole-blood samples of three PLWH individuals. CD3/28 was used to activate the CD4+ T cells for 3 days and then RNA was extracted for dRNA-seq. The significantly lower amount of HIV-1 viral RNA in these samples required longer sequencing. For one sample, patient 8, we also employed a method used by Baek and colleagues, inclusion of HIV-1-specific reverse transcription primers (Baek et al., 2024). However, this did not significantly boost coverage. Thus, only 3’ end transcript coverage was achievable for PLWH samples. For modification sites where we obtained sufficient coverage, we compared their levels to that of HIV-1 RNA isolated from Jurkat and CD4+ T cells grown in culture, as well as virion particles isolated from CD4+ T cell culture (**Figure 4C**). In general, modifications were found at the same positions and at similar frequencies compared to Jurkat cell culture samples. However, m^6^A modifications in cell-culture infected CD4+ T cells and virion samples were called at a lower frequency at nearly all positions. We noted that m^6^A at positions 8564, 8571, 8621, and 8710 were markedly higher for Jurkat cells as compared to other samples. The most likely explanation for this discrepancy is the presence of an altered sequence, specifically partial replacement of *nef* with GFP, in the NL4-3-GFP genome.

The conservation of m^6^A modifications, as well as the DRACH motifs where they occur, may hold functional significance. One consideration is the rate of mutation at these sites as compared to other nucleotide positions across the HIV-1 genome. HIV-1 is known to have a high mutation rate, one of the reasons why developing treatments is complicated (Yeo et al., 2020). The frequency of m^6^A modification was therefore correlated with the sequences covered in our PLWH samples and the Nextclade database of reference HIV-1 sequences (**Figure 4D**). Between HIV-1 genomic positions 8000 and 9172, there are twenty-six DRACH motifs. Of these, sixteen were not mutated in either sample. This includes nine high-frequency m^6^A modification sites. Of the ten DRACH motif sites that were disrupted in one or more PLWH samples, only A8710 showed a significant m^6^A modification frequency (47.7%) when extracted from Jurkat cell samples, but not in cultured CD4+ T cells and virions. We next compared against the Nextclade HIV-1 (HXB2) dataset, which comprises nucleotide diversity of approximately one thousand HIV-1 genomes (Aksamentov et al., 2021). For a given nucleotide position, one mutation observed for all the genomes is counted as one ‘event.’ A typical nucleotide position will have approximately fifty events. The highest number of mutation events at any nucleotide position was 406. In contrast, the lowest number of mutation events (1-5 events) are associated with essential genetic components, such as the start codons for HIV-1 proteins. When surveying the nine high-frequency m^6^A sites, the average number of events was 16.38, between 10-35% of the typical nucleotide mutation rate (**Figure 4D**). Together, these results suggest that these residues may be conserved for their m^6^A modification status and therefore play important roles in viral function and replication fitness.

## DISCUSSION

The role of chemical modifications to HIV-1 viral RNA in biology and disease, now referred to as epitranscriptomics, has been a growing topic of investigation for over three decades (Braddock et al., 1991; Sharmeen et al., 1991). Sequencing-based methodologies to both identify the types and map the locations of RNA modifications has significantly improved our understanding of their potential roles (Courtney, 2021; Imam et al., 2020; Phillips et al., 2024; Wang et al., 2022). These findings should enable sequence-specific manipulation of modifications for mechanistic studies as well as potential therapeutic strategies (Abudayyeh et al., 2019; Baek et al., 2024; Chen et al., 2019; Cox et al., 2017; Latifi et al., 2023; Reautschnig et al., 2024; Wilson et al., 2020; Xia et al., 2021; Zhang et al., 2024). Unfortunately, the diversity of sequencing-based methods, among other variables, has made it difficult to correlate findings across multiple studies or confidently probe the role of modifications. Standard inclusion of common experimental or bioinformatic controls would allow for straightforward normalization and correlation across many studies, providing a standard reference to benchmark against. Such a reference epitranscriptome could help ensure that results were reproducible, rigorous, and well-controlled and advance the study of HIV-1 epitranscriptomics by providing an initial level of ground truth.

Here, we sought to establish an HIV-1 reference epitranscriptome to strengthen data interpretation, better integrate results with those of other research groups, and create a foundation for HIV-1 epitranscriptomics to build upon. Commercial nanopore dRNA-seq reagents, workflows, and bioinformatic tools are broadly accessible and have undergone significant improvements. Thus, nanopore can provide a common tool and dRNA-seq results can offer a reproducible reference for detection of RNA modifications on HIV-1 viral RNAs. Existing data can also be reanalyzed when new algorithms become available, making it possible to map new modifications by simply repeating base-calling analyses or improve accuracy as algorithms evolve. In addition, current algorithms enable the equivalent of multiplexed modified base-calling, dramatically reducing resource demands. Together, these features make nanopore-based sequencing for epitranscriptomics cost-effective and data-rich (White and Hesselberth, 2022). When combined with commonly used reagents, such as widely accepted cell lines and genomes, a nanopore-based reference epitranscriptome becomes broadly useful.

We compared RNA preparation techniques, effects of cART, different cell lines, and virions, as well as characterized modifications in HIV-1 viral RNA samples from PLWH. We also constructed the first HIV-1 antisense epitranscriptome, which could provide unexpected insight into poorly understood HIV-1 antisense transcript biology. We found that nanopore-based dRNA-seq was generally robust and reproducible across replicates and sample and cell types. We discovered chemical modification features that have not been previously observed, such as the depletion of m^6^A but enrichment of inosine and psi on antisense HIV-1 viral RNAs, which appears to be the inverse for sense viral RNAs. We also discovered m^6^A and other modifications, albeit at a lower frequency, on HIV-1 viral RNAs from PLWH. These appear to be conserved when comparing NL4-3 to naturally circulating genomes, though their frequency at individual nucleotide positions may reflect cellular and environmental contexts. Interestingly, treatment with cART had no effect on modifications called at high confidence, suggesting that these drugs do not significantly alter the recognition and modification of HIV-1 RNAs.

Our results provide some implications for individual modifications in HIV-1 viral RNA. While the presence of m^5^C was previously reported by Cullen and colleagues using an antibody capture approach (Courtney et al., 2019), another group was unable to detect the same m^5^C by bisulfite sequencing methods (Huang et al., 2023). Our results suggest that m^5^C is present but at relatively low levels at only a handful of positions, possibly offering an explanation to the two conflicting reports. The m^5^C, inosine, and 2’-*O*-Me sites that remain at sufficiently high frequency after baseline correction may be worth further investigation. They are relatively distant from other modifications, suggesting they are not miscalled due to neighboring modifications, at least for those currently detectable by ONT algorithms. The high accuracy and precision of m^6^A calling we observed with ONT algorithms, especially at the 3’ end of the genome, combined with apparent conservation in naturally circulating HIV-1 strains, strengthens the functional significance of m^6^A in HIV-1 infection, host innate immunity, replication, and gene expression (Courtney, 2021; Imam et al., 2020; Phillips et al., 2024; Wang et al., 2022).

We observed differential modification calling frequencies on transcript splice isoforms. It is possible that splicing presents alternative accessibility to modifying enzymes due to RNA structure or protein interactions. Conversely, RNA modifications could alter splicing patterns by the same principles of altered RNA folding or protein binding. The potential order of events remains unclear, however, especially because some RNA modifying enzymes like METTL3 are known to be nuclear localized (Scholler et al., 2018). Nonetheless, the ability to detect splice isoform-specific modifications with nanopore will be useful in the study of HIV-1 regulatory mechanisms at the RNA or translation level, which help control the stoichiometry of viral factors (Emery and Swanstrom, 2021; Sertznig et al., 2018).

A limitation to this study, as well as nanopore-based epitranscriptomics more broadly, is the accuracy of algorithm-based modification calling (White and Hesselberth, 2022). We have demonstrated that identification of modifications can be influenced by both sequence context and neighboring modifications, which are difficult to fully account for. To partially address this, *in vitro* transcribed or chemically synthesized RNA fragments, with and without modifications, can be used to perform corrections and baseline subtractions. These analyses should be useful for better algorithm training, especially for specific, defined transcriptomes like HIV-1, and represent a general tool for establishing nanopore-based reference data sets. However, the effect of neighboring modifications on modification calling is not well understood and can affect the fidelity of identification in complex samples. Algorithms can only be expected to account for modifications and contexts they have been trained with. Similarly, new versions of algorithms, aimed at reducing noise, may lose sensitivity as a tradeoff for accuracy. During this study, ONT released the newest version of their RNA modification calling algorithm. We initially used the previous version to generate modification frequencies for all samples. The background of each modification under that algorithm (v5.1.0) (**Figure S6**) compared with the new algorithm (v5.2.0) (**Figure S3**) was significantly higher. However, the reduction in m^5^C and pseudouridine background calling with the new algorithm resulted in substantial loss of modifications at the baseline-corrected 10% threshold. Changes in significant modification calling (**Table S9**) exemplify the variation that can occur across algorithms. More sophisticated combinatorial data sets may be required to better train nanopore algorithms for calling RNA modifications in samples where multiple modifications are simultaneously present. Likewise, creating algorithms specific to the genome or transcriptome of a virus or organism, combined with synthetic modification contexts or mass spectrometry that defines the potential modification landscape, may represent a next step toward high-fidelity nanopore-based epitranscriptomics.

## METHODS

### Viral production and quantification

All HIV-1 isolates were generated by transfecting HEK 293T cells using polyethylenimine (PEI; PolySciences). Briefly, for a 10 cm plate production of VSV-G pseudotypes with GFP reporter genes, 6 µg of HIV-GFPΔEnv and 4 µg of pCMV-VSV-G (gifts from Dr. Thomas J. Hope) were combined as described previously (Mamede et al., 2017), and for replicative HIV-1 pNL4-3 we used 6 µg pNL4-3 (a gift from the HIV repository) with 1 mL of opti-MEM (Life Technologies). 40 µl of PEI was added to the mix, briefly vortexed, and allowed to incubate for 15 min before addition to HEK 293T cells. After overnight transfection, media was changed to fresh DMEM with 10% fetal bovine serum (FBS), 1x penicillin-streptomycin (P/S), and harvested at 48 h post-transfection through a 0.45 µm PVDF membrane filter. For replicative NL4-3, we concentrated the supernatants over a 20% sucrose cushion at 5600 x g overnight in 15 mL tubes. p24 ELISA was performed to quantify the viral preparations (R&D Systems, HIV-1 Gag p24 Quantikine ELISA Kit).

### Jurkat cell growth and infection

Jurkat cells (HIV repository) were cultured in RPMI-1640 medium supplemented with 10% FBS, 1x P/S, and non-essential amino acids (NEAA). Cells were split to 2 x 10^5^ cells/mL and split when they reached 1.5 x 10^6^ cells/mL. When infecting with replicative NL4-3 and pHIVdEnv-GFP, we used a high cell density of ∼2.5 x 10^6^ cells/mL with a working concentration of HIV p24 (determined by ELISA) of 500-600 ng/mL. Cells were harvested at 45-48 h post-infection and TRIzol (Invitrogen) added at 1 mL per 1 x 10^7^ cells.

### CD4+ T cell negative selection, activation, growth, and infection

In whole blood tubes from unidentified healthy donors or leukopaks (Red Cross Blood) we performed negative cell separation following the vendors protocol with EasySep Direct Human CD4+ T cell Isolation Kits or RosetteSep Human CD4+ T cell Enrichment Cocktail (Stem Cell Technologies) with Lymphoprep (Stem Cell Technologies) as density medium, respectively. Before infection, we activated T cells by adding anti-CD3 and anti-CD28 antibodies along with IL-2 following the methods described previously (Mamede et al., 2017). Activation was confirmed 3 d post-treatment by measuring CD4+/CD69+ double-positive cell percentage by flow cytometry. Briefly: Day 0, isolate CD4+ T cells from blood and activate T cells; Day 3: test activation % and infect CD4+ T cells with WT-HIV. Day 4: TRIUMEQ cART treatment when applicable; Day 6: harvest CD4+ T cells for TRIzol lysis as well as quantify infection rate by flow cytometry with KC57 antibody (as described below).

### Patient sample CD4+ T cell collection, preparation, and latency reversal

Approximately 10 tubes of whole blood were collected in heparin tubes typically totaling 85 mL of whole blood. CD4+ T cells were negatively selected with RosetteSep Human CD4+ T cell Enrichment Cocktail (Stem Cell Technologies) and SepMate tubes with Lymphoprep (Stem Cell Technologies) as density medium following the vendor protocol. Cells were activated with anti-CD3 and anti-CD28 antibodies as described above for 3 d. On day 3 cells were fixed for flow cytometry to quantify the percentage of positive p24 cells as described below. The serum on top of the gradient media after centrifugation was collected. Viral particles were concentrated by ultracentrifugation with a 20% sucrose cushion in PBS for 4 h at ∼120,000 x g using an SW28 rotor (Beckman Coulter). After concentration, the supernatant was discarded and TRIzol was added to the viral pellets (2 mL total for the initial ∼85 mL of plasma). Patient sample collections were approved by the Institutional Review Board (IRB) of Rush University (FWA number: 00000482; ORA Number: 21072211-IRB01).

### Drug treatments

To model current cART therapy, we added TRIUMEQ with the respective concentrations for each of the antiretrovirals based on reported plasma levels: *ABC 140 ng/mL, 3TC 670 ng/mL, DTG 3.9 ug/mL* (Best et al., 2011; Letendre et al., 2014; McDowell et al., 1999; Mueller et al., 1998; Tashima et al., 1999; van Praag et al., 2002). For STM2457 (MedChemExpress) treatment, stocks were dissolved in dimethyl sulfoxide (DMSO) and the drug was added simultaneously to infection of NL4-3 at a final of 30 µM.

### Flow cytometry

Flow cytometry was performed with BD CytoFix/Cytoperm per vendor protocol with the following antibodies: to measure activation, anti-CD69 (Fisher Scientific BDB560968) and anti-CD25; to measure Gag/p24 in the cytoplasm to report infection or viral production, KC57-RD1 (Beckman Coulter, Cat#6604667) antibody (or isotype). Samples were analyzed in a LSR Fortessa (BD Biosciences) gating for doublets with SSC-H/SSC-W and FSC-H/FSC-W gates.

### RNA extraction and preparation

Total RNA was extracted from cell pellets using the manufacturer’s recommended TRIzol (Invitrogen) protocol. Ribosomal RNA was removed by enzymatic rRNA depletion or poly-A RNA was selected by bead-based enrichment. For rRNA depletion, approximately 20 µg of total RNA was hybridized with DNA oligonucleotides (**Table S10**) complementary to rRNA (Baldwin et al., 2021). The solution was heated to 95°C and slow-cooled to 65°C to enable oligonucleotide hybridization. RNase H (McLabs, HTRH-200) was added (200 U) and the solution incubated at 65°C for 5 min to degrade rRNA. DNA was then degraded using 150 U of Turbo DNase (Invitrogen, AM2239) at 37°C for 20 min. RNA was then purified using AMPure XP beads, typically yielding approximately 1000 ng of recovered RNA.

For poly-A selection, the NEBNext High Input Poly(A) mRNA Isolation Module (E3370S) was used following the manufacturer’s recommended protocol. Briefly, approximately 20 µg of total RNA was added to a solution containing poly-A-binding beads. After several wash steps, the bead-bound RNA is eluted by heating, typically yielding approximately 500 ng of recovered RNA.

### Synthetic and *in vitro* transcribed HIV-1 RNA fragments

For small unmodified and m^6^A-modified RNA fragments, 60 nucleotide RNAs were prepared by solid-phase chemical synthesis, HPLC purification, and mass spectrometry confirmation following previously published methods (Ageely et al., 2021; Lackey et al., 2009). Commercially available m^6^A phosphoramidite was used (Glen Research). For *in vitro* transcription, pNL4-3 HIV-1 sequence was used to create primers for overlapping amplicons that span the HIV-1 genome (**Table S10**). Primers were created with or without a 5’ T7 promoter sequence. PCR amplification resulted in T7 promoter-containing fragments of the genome of 2-3 kb size, which were then used as templates in T7 RNA polymerase transcription reactions to make either sense or antisense RNA HIV-1 fragments. Briefly, approximately 1 µg of template DNA was used in a 60 µL reaction following previously published conditions and enzyme preparations (Kartje et al., 2021). For transcripts containing modified nucleotides, native ribonucleotide triphosphates (NTPs) were fully replaced with modified variants from TriLink Biotechnologies, either N^6^-methyladenosine-5’-triphosphate (N-1013) for ATP, 5-methylcytidine-5’-triphosphate (N-1014) for CTP, or pseudouridine-5’-triphosphate (N-1019) for UTP. Reactions were incubated at 37°C for 90 min then treated with DNase I for an additional 20 min at 37°C. RNA was isolated by phenol/chloroform extraction and ethanol precipitation. The resulting RNA fragments were combined and sequenced by nanopore in the same manner as cell-derived RNA.

### Nanopore direct RNA sequencing

For rRNA-depleted samples, polyadenylation reactions were performed using *E. coli* Poly(A) Polymerase (New England Biolabs, M0276L). Briefly, 1000 ng of rRNA-depleted RNA was treated with 0.375 U/µL *E. coli* Poly(A) Polymerase in 1x *E. coli* Poly(A) Polymerase Buffer and 1 mM ATP. The reaction was incubated at 37°C for 2 min and stopped by addition of EDTA to a final concentration of 10 mM. The reaction was bead-cleaned using 2x RNAClean XP Beads (Beckman Coulter), washed twice with 200 µL of 75% ethanol, and eluted in 7 µL of nuclease-free water. Concentration was determined using the Qubit RNA HS Kit (ThermoFisher Scientific).

Library preparation of the RNA samples was completed using the direct RNA sequencing kit (SQK-RNA004; Oxford Nanopore Technologies, ONT), following manufacturer’s recommended protocol. Briefly, RNA samples were quantified using Qubit RNA HS Kit (ThermoFisher Scientific). 1 µL SUPERase·In was added, as well as 1 µL of 130 µM pooled reverse transcription (RT) primers reported by Baek and colleagues (Baek et al., 2024) for sample 13B (patient 8 CD4+ T cell) and 14H (CD4+ virions), and the mixture heated at 65°C for 5 min then cooled to 4°C. The RT Adapter was ligated by adding 3 μL of 5x Quick Ligation Buffer (New England Biolabs, NEB), 2 μL of T4 DNA ligase at 2000 U/μL (NEB), 1 μL SUPERase·In RNase Inhibitor (20 U/μL) (ThermoFisher Scientific) and incubated at room temperature for 15 min. Reverse transcription was performed using a master mix containing 1x First-Strand Buffer, 10 mM DTT, 0.5 mM dNTPs, and nuclease-free water. The master mix was added to the RT adapter-ligated RNA, followed by the addition of SuperScript III Reverse Transcriptase (ThermoFisher Scientific). Reactions were incubated in a thermal cycler at 50°C for 60 minutes, and 70°C for 10 minutes, with a final hold at 4°C. The reaction was purified using 1.8x RNAClean XP beads (Beckman Coulter), washed twice with 75% ethanol and eluted in 20 µL nuclease-free water. RNA adapter (RMX) ligation was performed by mixing 20 µL of the reverse transcribed RNA, 1x NEBNext Quick Ligation Buffer, 6 µL RNA Adapter (RLA), 3 µL nuclease-free water, and 3 µL T4 DNA Ligase at 2000 U/μL (NEB). The mixture was incubated at room temperature for 15 minutes. The reaction was purified using 0.4x RNAClean XP beads (Beckman Coulter), washed twice with Wash Buffer (WSH) and eluted in 33 µL of RNA elution buffer (REB). The final library was loaded on a PromethION RNA flow cell (FLO-PRO004RA) and sequenced on a PromethION P2 solo using MinKNOW (v. 24.06.10) with POD5 and live base-calling on. Coverage was monitored using RAMPART software (https://artic.network/rampart), with a goal of 50x coverage across the HIV-1 genome.

### Modification calling from Nanopore POD5 files

After the completion of sequencing, reads were called from the POD5 files with Dorado software (v1.0.0) with settings for RNA modifications (RNA basecalling model rna004_130bps_sup@v5.2.0, https://github.com/nanoporetech/dorado). Generated reads were automatically aligned against a FASTA file containing human (hg38) and HIV-1 genomes (https://www.ncbi.nlm.nih.gov/nuccore/AF324493), generating a BAM file.

Dorado basecaller sup,m5C_2OmeC,inosine_m6A_2OmeA,pseU_2OmeU,2OmeG [pod5_folder_location] -b 408 --min-qscore 8 --device cuda:all –mm2-opts “-x splice” --reference [reference_fasta_location] > unsorted_reads.bam

After generation of the BAM file, samtools was used to sort and index the BAM file.

~~~
samtools sort unsorted_reads.bam > sorted_reads.bam
samtools index sorted_reads.bam
~~~

This sorted BAM file was then processed by modkit (https://github.com/nanoporetech/modkit) to generate a file containing modification sites on the HIV-1 genome. A filter-threshold of 0.7 was utilized to standardize results between runs.

`modkit pileup [location_of_BAM] --ref [loc_of_reference] -t 18 -- region NL43_AF324493.2_R2R --max-depth 10000000 --bedgraph ./ --filter-threshold .7 --motif DRACH 2 --prefix m6A`

`modkit pileup [location_of_BAM] --ref [loc_of_reference] -t 18 -- region NL43_AF324493.2_R2R --max-depth 10000000 --bedgraph ./ --filter-threshold .7 --motif C 0 --prefix m5C`

`modkit pileup [location_of_BAM] --ref [loc_of_reference] -t 18 -- region NL43_AF324493.2_R2R --max-depth 10000000 --bedgraph ./ --filter-threshold .7 --motif T 0 --prefix Psi`

`modkit pileup [location_of_BAM] --ref [loc_of_reference] -t 18 -- region NL43_AF324493.2_R2R --max-depth 10000000 --bedgraph ./ --filter-threshold .7 --motif A 0 --prefix Ino`

The four 2’-*O*-methylated bases can be similarly called with motifs ‘A 0’, ‘T 0’, ‘C 0’ and ‘G 0’. These modkit commands will generate multiple files per command, with the filename differentiating between the sense and antisense direction of the reference, and between the modification being identified, with each modification having an identification code:

| Modification | modkit ID |
| --- | --- |
| m<sup>6</sup>A | a |
| m<sup>5</sup>C | m |
| pseudouridine | 17802 |
| inosine | 17596 |
| A<sub>m</sub> | 69426 |
| U<sub>m</sub> | 19227 |
| C<sub>m</sub> | 19228 |
| G<sub>m</sub> | 19229 |

The result is a bedgraph file containing the reference genome name, the position of the calculated modification, the percentage of reads that had the modification, and the total number of reads in the BAM file at that nucleotide position, e.g.:

~~~
NL43_AF324493.2_R2R       8988   8989   0.64912283   57
~~~

To remove noise from low-coverage nucleotides, a filter was applied to the bedgraph file, removing modification locations that had less than 10x coverage for that nucleotide position.

~~~
awk ‘$5>9’ m6A_original.bedgraph > m6A_10xFilter.bedgraph
~~~

The resulting bedgraph file can be viewed in IGV (https://igv.org/) or opened directly in text-editing software as a tab-separated value (TSV) file.

### Baseline correction of modification calling

RNA fragments *in vitro* transcribed and sequenced as described above were modification called the same as other samples. RNA fragments *in vitro* transcribed without modified bases were used as a baseline for all other samples since any modification calls would be considered false. For each RNA sample used in the study, there is a modification frequency for each individual modification at each individual nucleotide that this modification could be present at. To correct sequence pattern-based miscalls, the T7 *in vitro* transcribed values for a particular modification and nucleotide position were subtracted from the value found for a sample at each nucleotide position. If this subtraction brought the value below zero, zero was used instead of a negative value.

### Separating splice isoforms using FLAIR

Full-Length Alternative Isoform analysis of RNA (FLAIR) software was used to identify the splice isoforms of HIV-1 reads and generate readID lists of these isoforms (Tang et al., 2020). These lists were used to separate splice isoforms into separate BAM files that could be called for modifications individually.

Full scripts and reference files can be found at https://github.com/gagnonlab/HIV-dRNA-Seq. Briefly, the aligned BAM file generated previously from modification calling is processed by FLAIR to identify the splice isoform of as many reads as possible. If a read could fit multiple isoforms, it is excluded.

`flair quantify -r manifest.tsv -i hiv_isoforms.fa --output quantify/_--threads 18 --generate_map --stringent --isoform_bed hiv_isoforms.bed`

manifest.tsv is a list of BAM files to be processed. hiv_isoforms.fa is a FASTA file of each of the splice isoforms, based on the NL4-3 HIV-1 genome with the added GFP tag. hiv_isoforms.bed is a BED file mapping the different splice isoforms. Both are generated from a GTF file containing the splice isoforms. This process generates a read.map.txt file that contains a list of each of the input splice isoforms, followed by a comma-separated listing of all readIDs that match that splice isoform.

The list of readIDs is processed using a bash script to separate out these listed readIDs from a BAM file into individual BAM files for each splice isoform. The full script can be found on the lab GitHub, https://github.com/gagnonlab/HIV-dRNA-Seq. Briefly, it converts the binary BAM file into a text-readable SAM file, copies the header of the SAM file into each individual splice file, uses the grep command to match reads from the full SAM file to readIDs from each splice isoform, and then copies these reads into the appropriate SAM output file. The output SAM files are converted back into BAM files and indexed. Afterwards, the resulting BAMs can be modification-called using Dorado as above.

## ACKNOWLEDGEMENTS

This project was funded by a National Institutes of Allergy and Infectious Diseases grant (5R61AI169661) to K.T.G. and J.I.M.

## DATA ACCESSIBILITY

Raw nanopore read data (.POD5), basecalled data (.BAM), lists of modification positions and frequency (.bedgraph), and genomic reference files and bash scripts reported in this manuscript are available at https://zenodo.org/record/15596052 (DOI: 10.5281/zenodo.15596052). Files are compressed together into a single .tar.xz file per sample.

---

**Figure S1.**
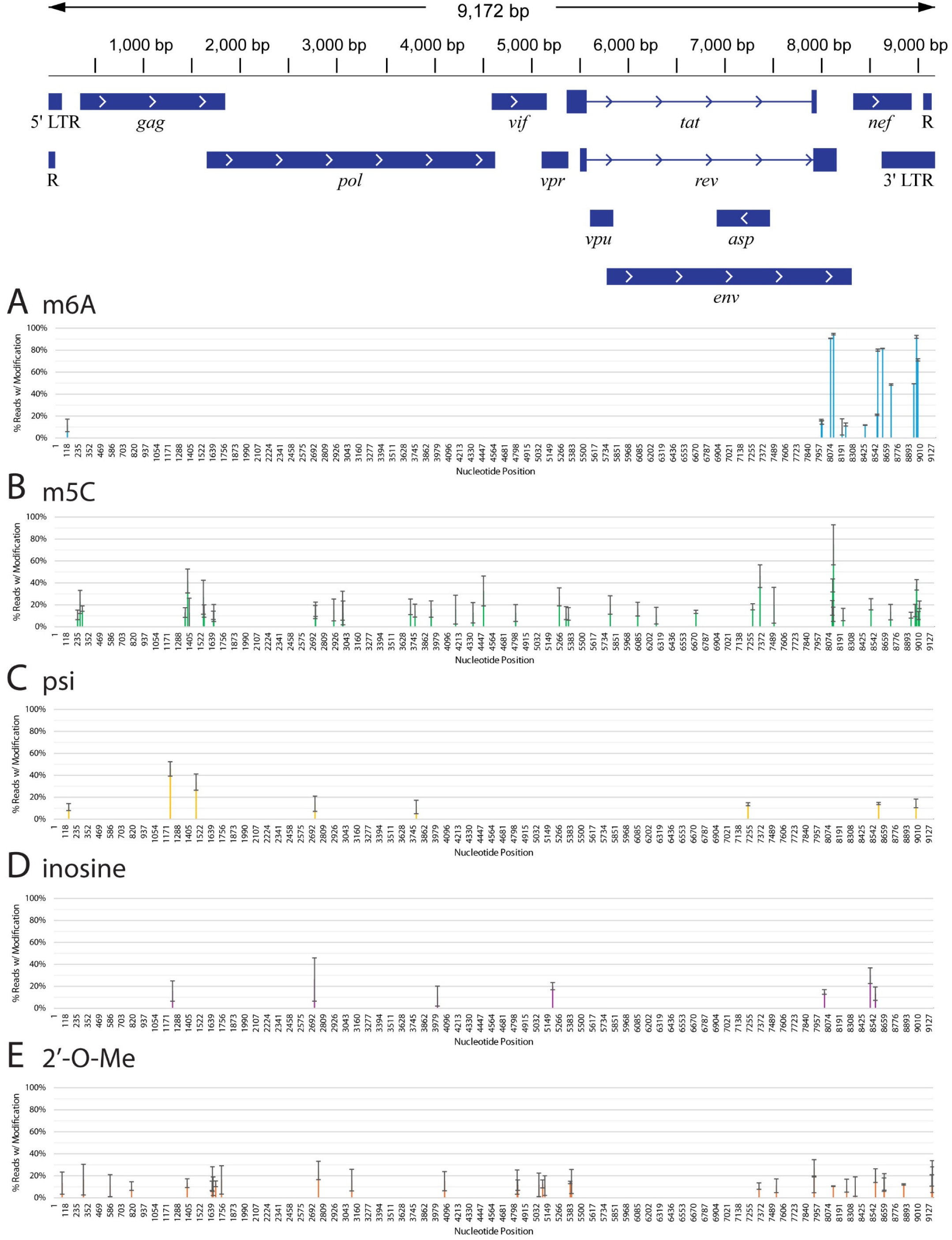
Uncorrected nanopore modification-calling results for HIV-1 viral RNA from Jurkat cells. X-axis represents individual nucleotides in the HIV-1 genome. Y-axis represents percentage of reads that contained the mutation of interest for that nucleotide. m^6^A (blue), m^5^C (green), pseudouridine (psi) (yellow), inosine (purple), and 2’-*O*-methylation (orange). Results are the average of three separate biological replicates. Error bars are standard deviation. Only results with percentages above 10% are shown.

**Figure S2.**
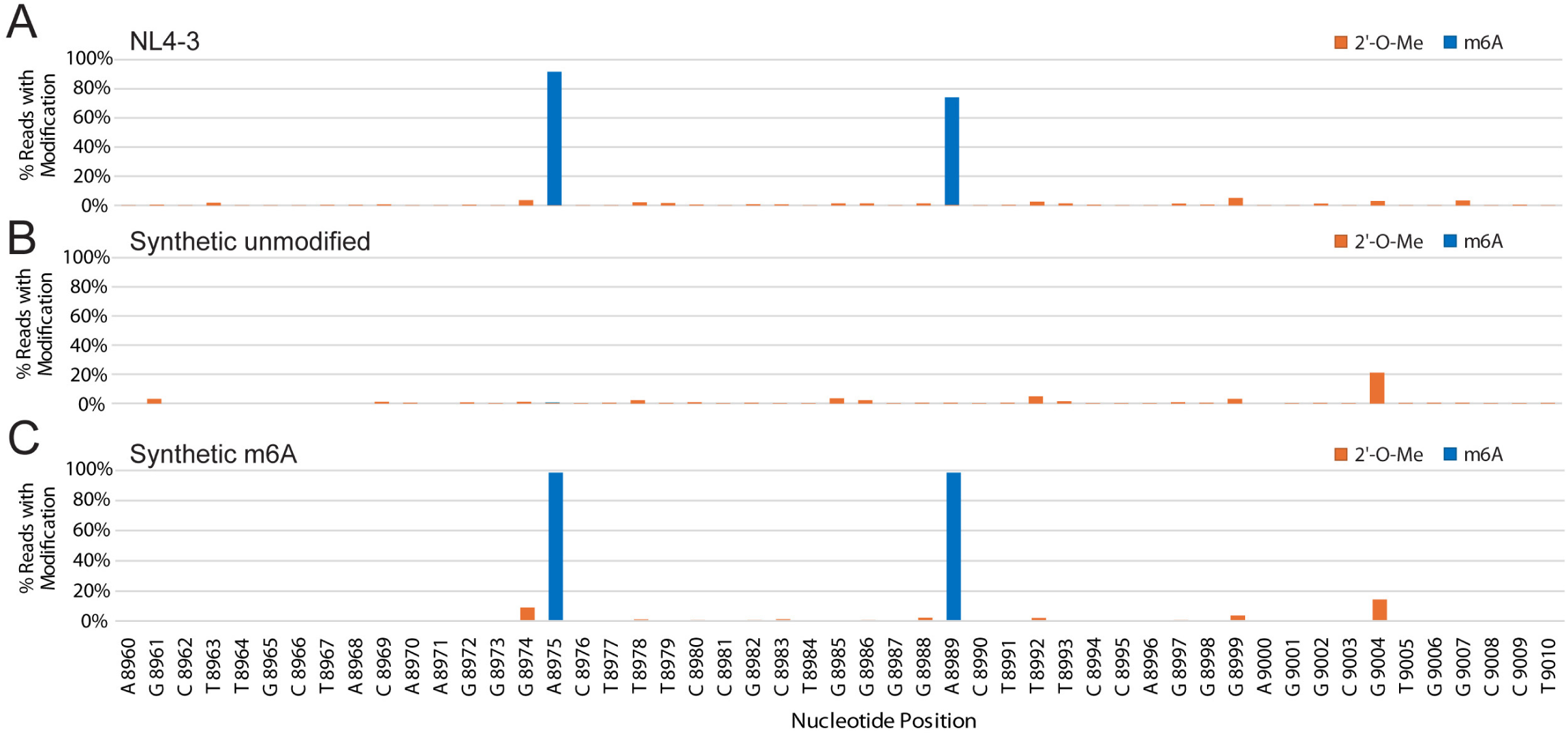
2’-*O*-methylation calling in HIV-1 RNA and synthetic RNA fragments. Comparison of modification calling between NL4-3 from Jurkat cells (upper) and two synthetic HIV-1 RNA fragments, one unmodified (middle) and one bearing m^6^A (lower) at two DRACH motifs. The nucleotide position corresponding to the NL4-3 genome is indicated on the x-axis. All modifications called in middle panel are incorrect while modifications called in the lower panel, besides m^6^A at position 8975 and 8989, are incorrect.

**Figure S3.**
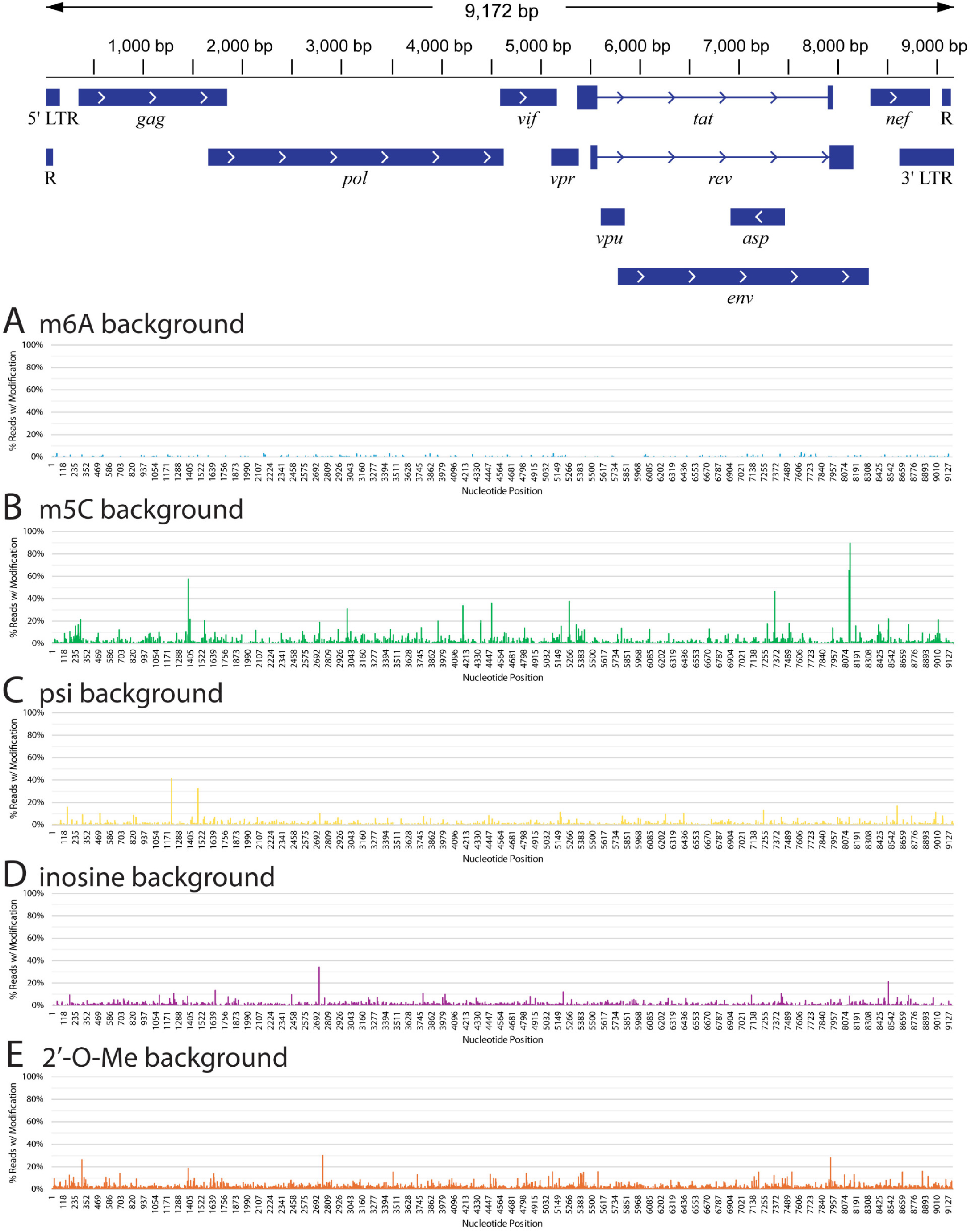
Modification calling from nanopore sequencing of *in vitro* transcribed NL4-3 HIV-1 RNA fragments used for baseline correction. X-axis represents individual nucleotides in the HIV-1 genome. Y-axis represents percentage of reads that contained the mutation of interest for that nucleotide. m^6^A (blue), m^5^C (green), pseudouridine (psi) (yellow), inosine (purple) and 2’-*O*-methylation (orange).

**Figure S4.**
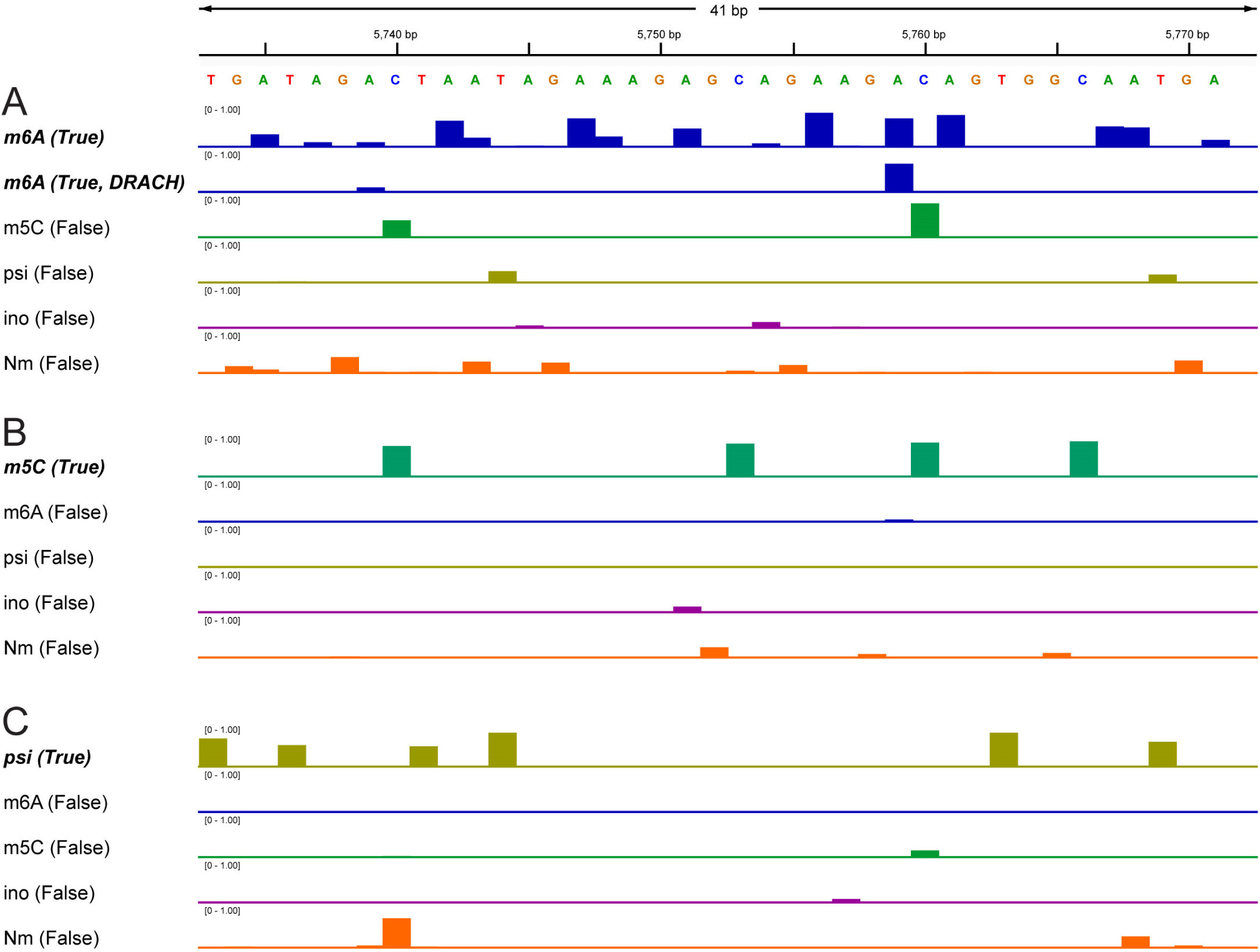
Miscalled modifications due to adjacent modifications. X-axis of each panel is the nucleotide position that corresponds to the sequence at the top of the figure. Y-axis is the percentage of reads containing a modification that position, according to Nanopore RNA modification-calling algorithm. m^6^A (blue), m^5^C (green), psi (yellow), inosine (purple), and 2’-*O*-Me (orange). (**A**) is derived from t7 *in vitro* transcribed RNA in which all As have been replaced with m6A. (**B**) is derived from t7 *in vitro* transcribed RNA in which all Cs have been replaced with m5C. (**C**) is derived from t7 *in vitro* transcribed RNA in which all Us have been replaced with psi.

**Figure S5.**
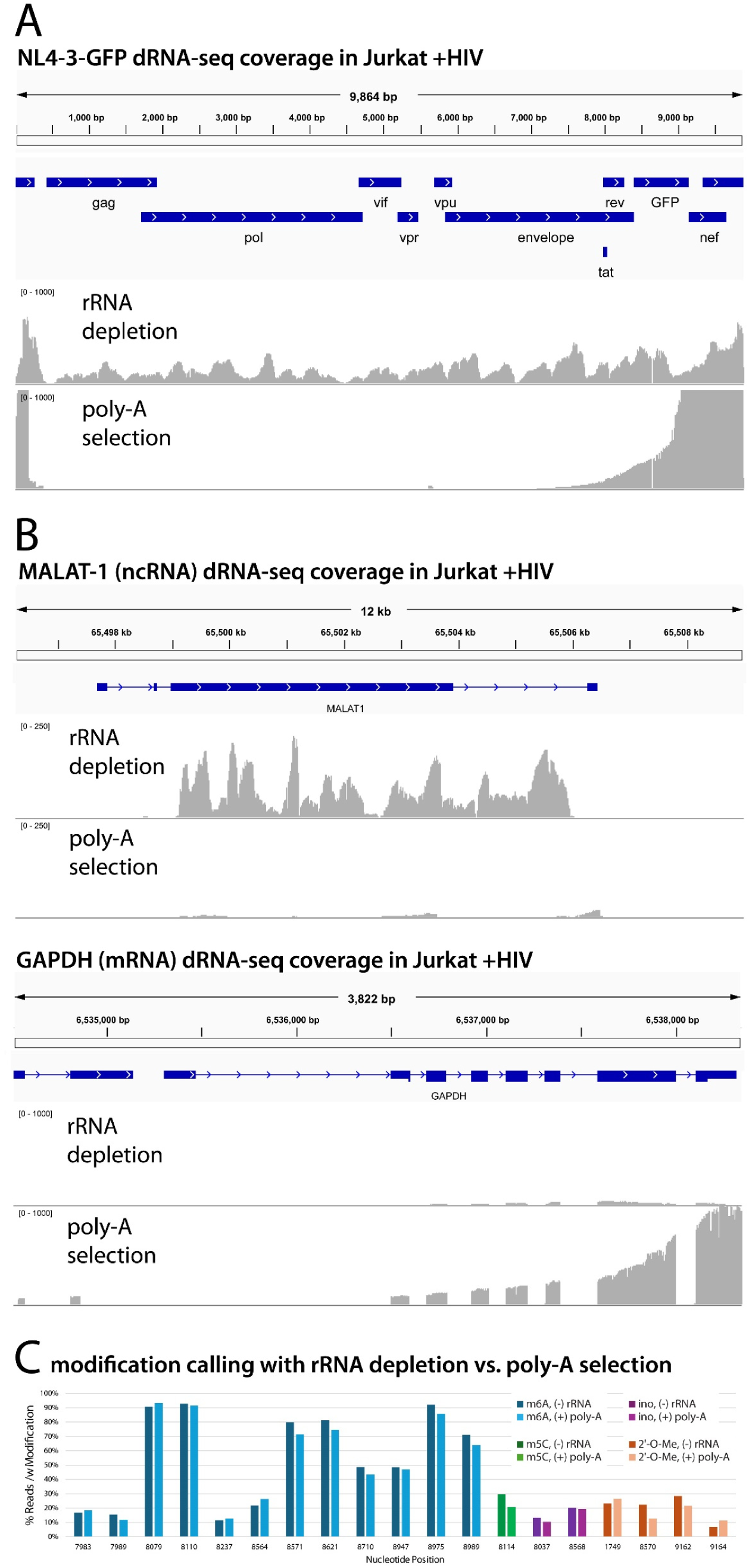
Comparison of rRNA depletion and poly-A selection using read coverage maps and modified base calling from nanopore dRNA-seq. X-axis is the individual nucleotide positions. Y-axis is the number of reads with coverage at that position. Samples are from Jurkat cells infected with NL4-3-GFP HIV-1. (**A**) Comparison of HIV-1 sequencing coverage from samples prepared by rRNA depletion or poly-A selection. (**B**) Comparison of human noncoding RNA MALAT-1 and mRNA GAPDH sequencing coverage for each RNA preparation method. (**C**) Modification calling results for HIV-1 viral RNA taken from Jurkat cell cultures and rRNA depleted or poly-A selected prior to dRNA-seq. m^6^A (blue), m^5^C (green), pseudouridine (psi) (yellow), inosine (purple) and 2’-*O*-methylation (orange).

**Figure S6.**
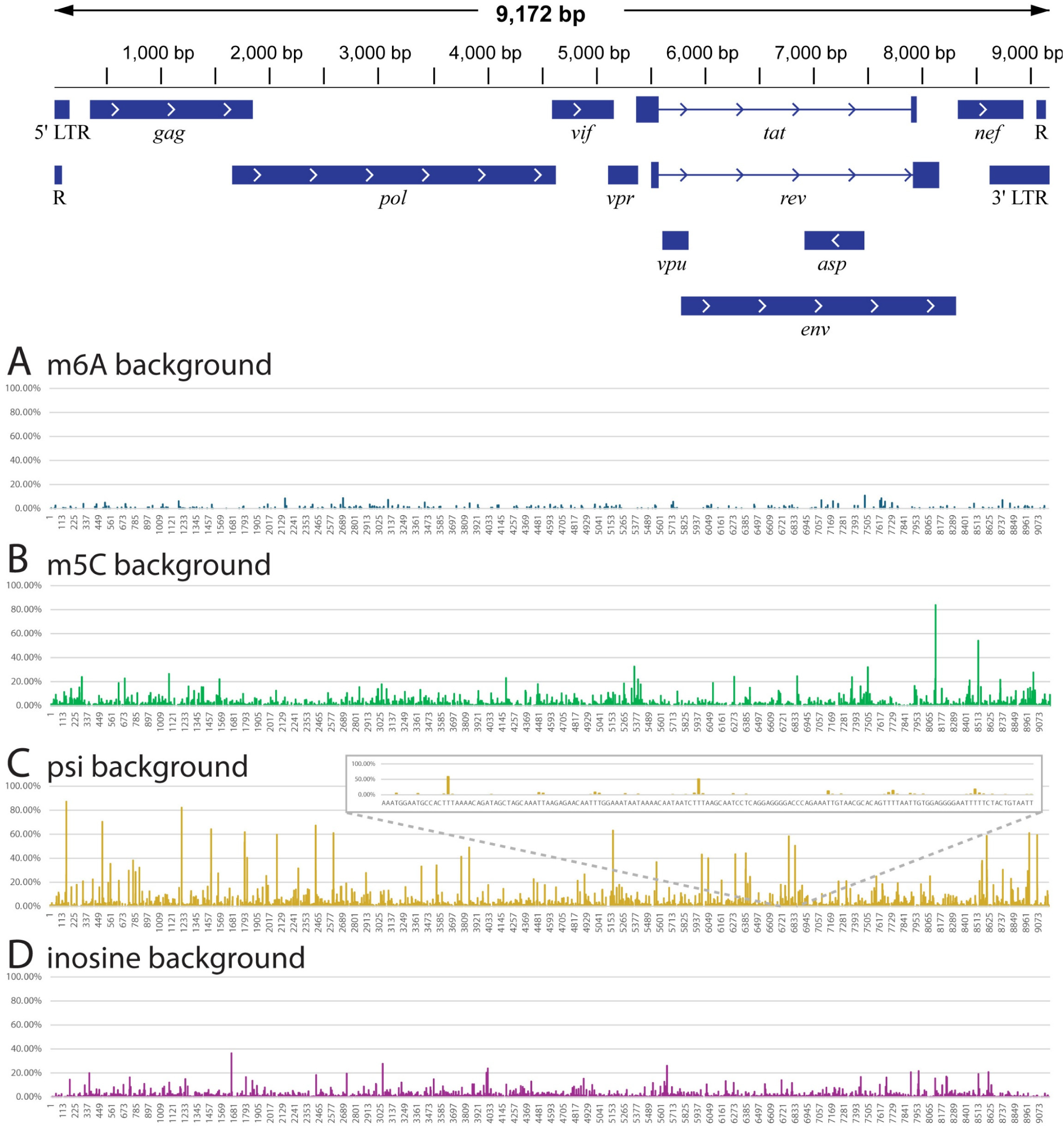
Modification calling from nanopore sequencing of *in vitro* transcribed NL4-3 HIV-1 RNA fragments used for baseline correction, using older Nanopore modification calling algorithm. Utilizes Dorado RNA modification basecalling algorithm rna004_130bps_sup@v5.1.0. X-axis represents individual nucleotides in the HIV-1 genome. Y-axis represents percentage of reads that contained the mutation of interest for that nucleotide. m^6^A (blue), m^5^C (green), pseudouridine (psi) (yellow), inosine (purple). Dorado algorithm rna004_130bps_sup@v5.1.0 does not support 2’-*O*-methylation calling.

**Table S1.**
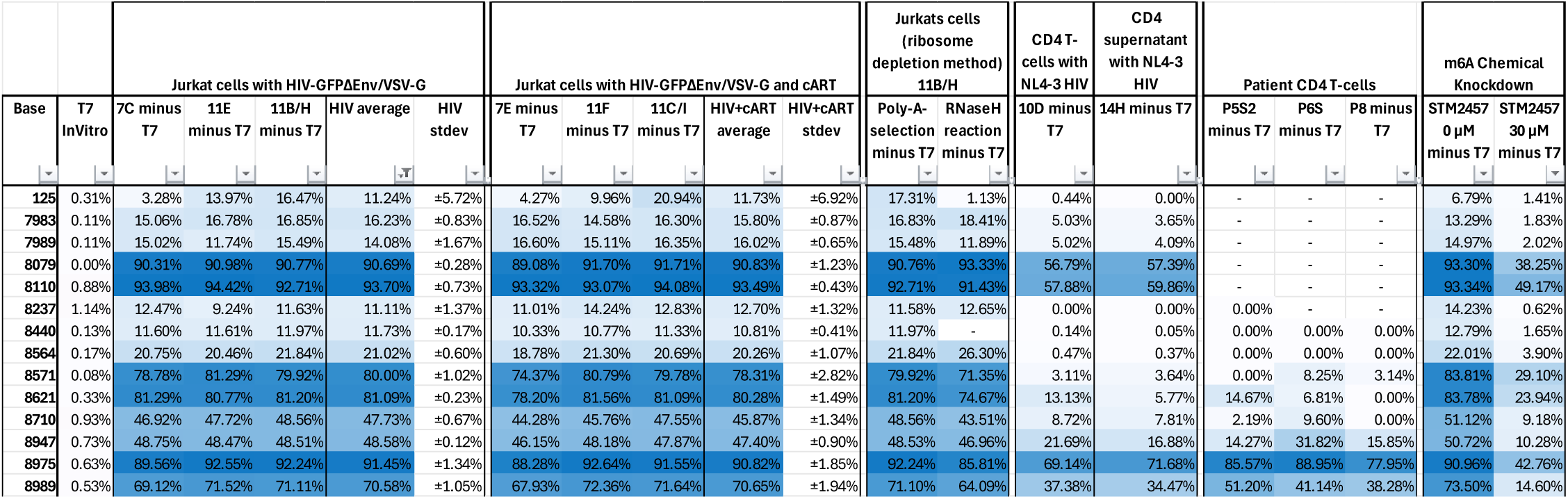
Significant m6A modification sites in the HIV-1 sense epitranscriptome.

**Table S2.**
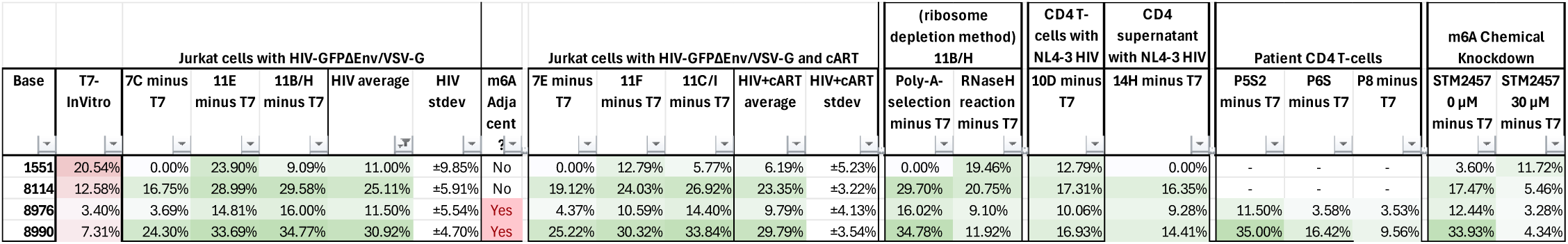
Significant m5C modification sites in the HIV-1 sense epitranscriptome.

**Table S3.**
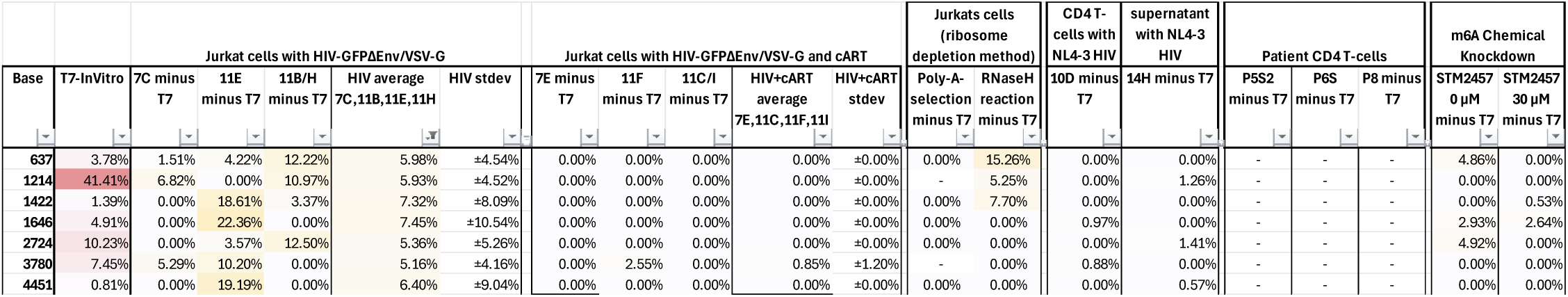
Significant pseudouridine modification sites in HIV-1 sense epitranscriptome (low threshold).

**Table S4.**
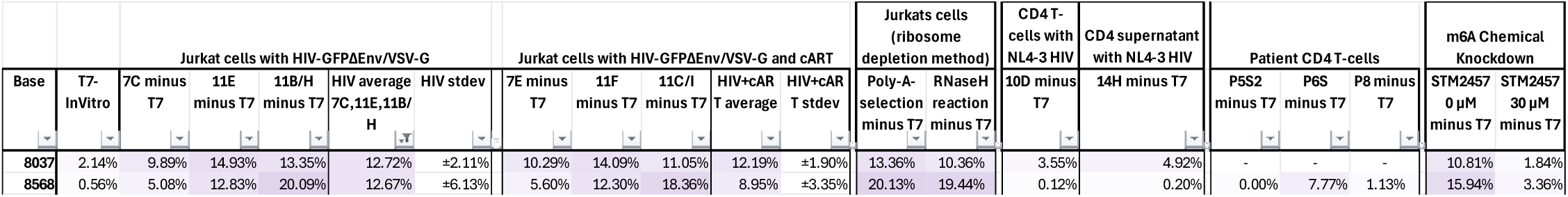
Significant inosine modification sites in the HIV-1 sense epitranscriptome.

**Table S5.** Significant 2’-O-methylation modification sites in the HIV-1 sense epitranscriptome.

|  |  | Jurkat cells with HIV-GFPΔEnv/VSV-G |  |  |  |  | Jurkat cells with HIV-GFPΔEnv/VSV-G and cART |  |  |  |  | Jurkats cells (ribosome depletion method) 11B/H |  | CD4 T-cells with NL4-3 HIV | CD4 supernatant with NL4-3 HIV | Patient CD4 T-cells |  |  | m6A Chemical Knockdown |  |
| --- | --- | --- | --- | --- | --- | --- | --- | --- | --- | --- | --- | --- | --- | --- | --- | --- | --- | --- | --- | --- |
| Base | T7 InVitro | 7C minus T7 | 11E minus T7 | 11B/H minus T7 | HIV average | HIV stdev | 7E minus T7 | 11F minus T7 | 11C/I minus T7 | HIV+cART average | HIV+cART stdev | Poly-A selection minus T7 | RNaseH reaction minus T7 | 10D minus T7 | 14H minus T7 | P5S2 minus T7 | P6S minus T7 | P8 minus T7 | STM2457 0 μM minus T7 | STM2457 30 μM minus T7 |
| 1749 | 6.82% | 0.00% | 9.85% | 25.00% | 11.62% | ±10.28% | 0.00% | 0.00% | 7.47% | 2.49% | ±3.52% | 23.18% | 26.52% | 5.68% | 0.00% | - | - | - | 9.46% | 0.00% |
| 3107 | 4.72% | 21.18% | - | 1.53% | 11.36% | ±9.82% | 16.15% | 0.00% | 0.00% | 5.38% | ±7.61% | 0.00% | 0.00% | 0.00% | 0.00% | - | - | - | 0.00% | 0.00% |
| 9162 | 7.41% | 12.96% | 18.81% | 28.46% | 20.08% | ±6.39% | 18.32% | 23.16% | 25.52% | 22.33% | ±3.00% | 28.47% | 21.53% | 20.02% | 26.28% | 41.42% | 31.16% | 26.83% | 23.57% | 24.46% |
| 9164 | 1.27% | 31.69% | 6.97% | 7.00% | 15.22% | ±11.65% | 26.43% | 13.06% | 5.20% | 14.90% | ±8.76% | 7.00% | 11.32% | 4.83% | 6.08% | 0.29% | 6.96% | 12.08% | 6.60% | 6.56% |

**Table S6.**
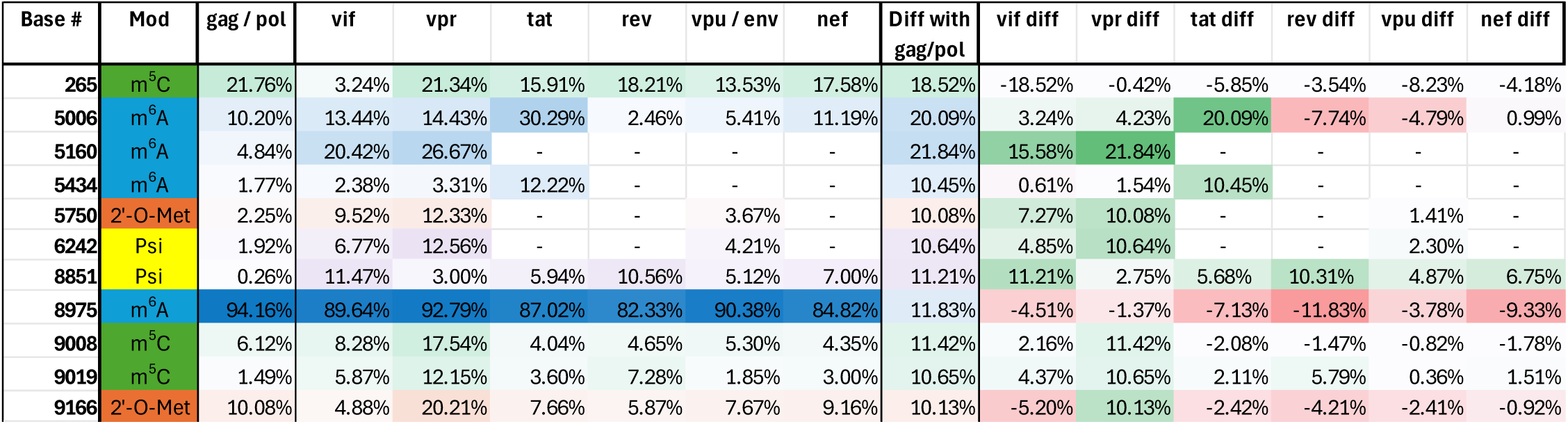
Significant differential modifications in transcript splice isoforms.

**Table S7.**
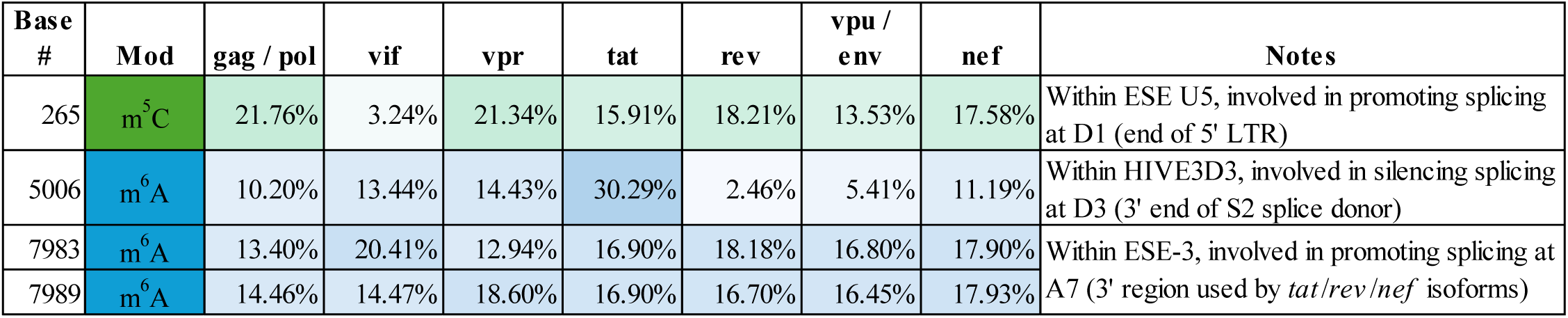
Modifications that fall within a known splicing regulatory element.

**Table S8.**
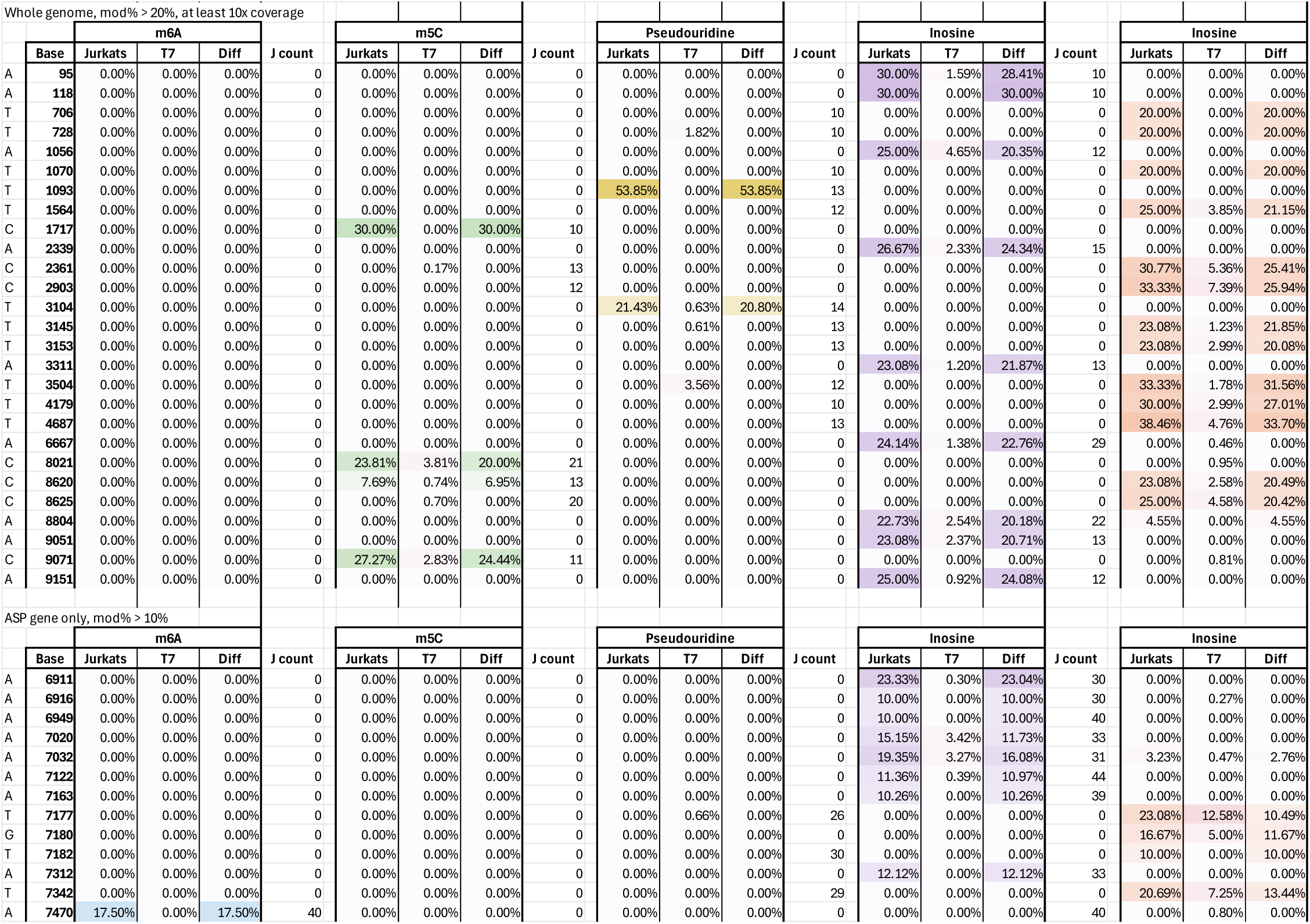
Significant modification sites in the NL4-3-GFP HIV-1 antisense epitranscriptome.

**Table S9.**
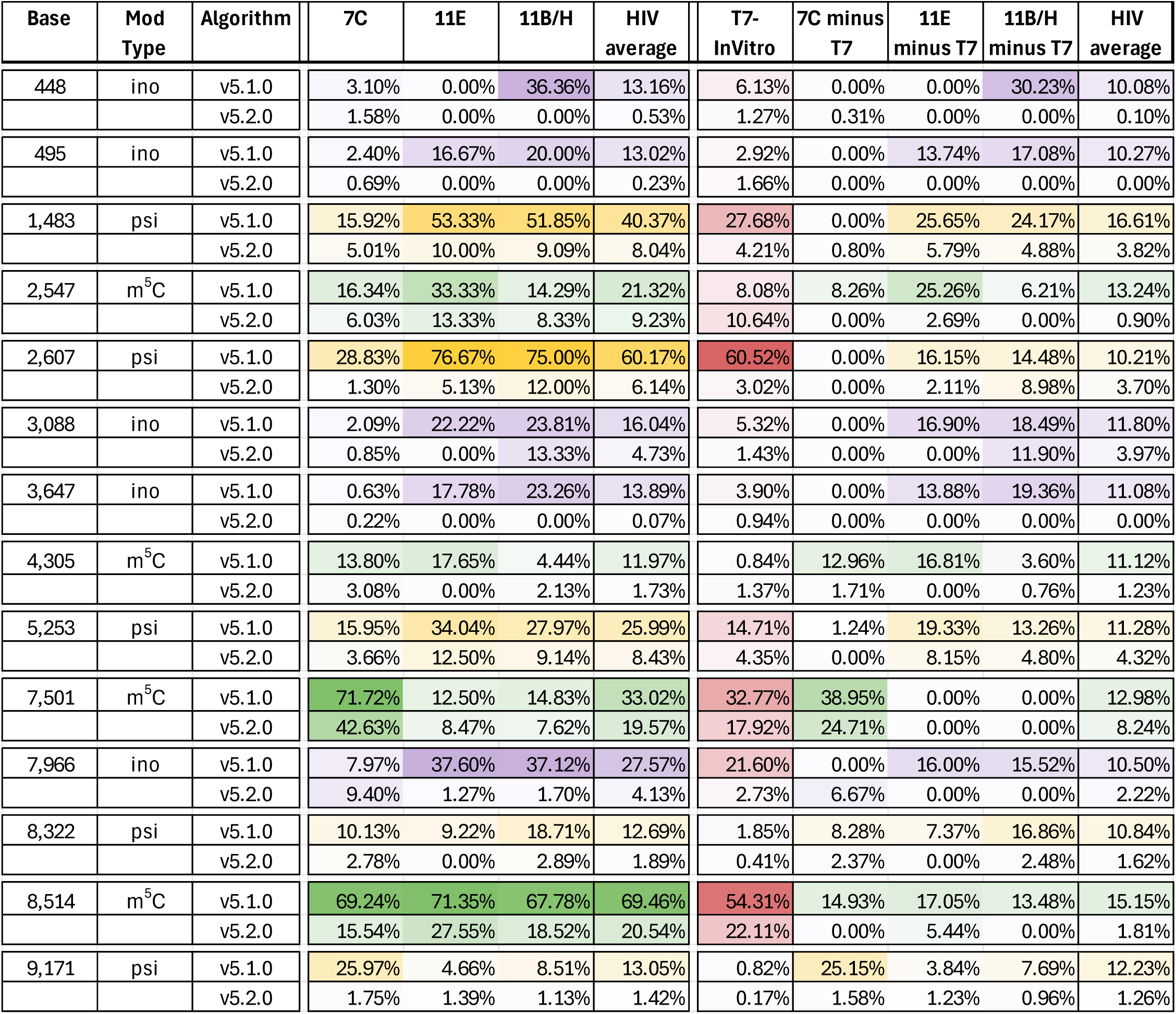
Comparison of modification frequencies between new (v5.2.0) and old (v5.1.0) Oxford Nanopore Technologies modification-calling algorithms.

| Base | Mod Type | Algorithm | 7C | 11E | 11B/H | HIV average | T7-InVitro | 7C minus T7 | 11E minus T7 | 11B/H minus T7 | HIV average |
| --- | --- | --- | --- | --- | --- | --- | --- | --- | --- | --- | --- |
| 448 | ino | v5.1.0 | 3.10% | 0.00% | 36.36% | 13.16% | 6.13% | 0.00% | 0.00% | 30.23% | 10.08% |
|  |  | v5.2.0 | 1.58% | 0.00% | 0.00% | 0.53% | 1.27% | 0.31% | 0.00% | 0.00% | 0.10% |
| 495 | ino | v5.1.0 | 2.40% | 16.67% | 20.00% | 13.02% | 2.92% | 0.00% | 13.74% | 17.08% | 10.27% |
|  |  | v5.2.0 | 0.69% | 0.00% | 0.00% | 0.23% | 1.66% | 0.00% | 0.00% | 0.00% | 0.00% |
| 1,483 | psi | v5.1.0 | 15.92% | 53.33% | 51.85% | 40.37% | 27.68% | 0.00% | 25.65% | 24.17% | 16.61% |
|  |  | v5.2.0 | 5.01% | 10.00% | 9.09% | 8.04% | 4.21% | 0.80% | 5.79% | 4.88% | 3.82% |
| 2,547 | m<sup>5</sup>C | v5.1.0 | 16.34% | 33.33% | 14.29% | 21.32% | 8.08% | 8.26% | 25.26% | 6.21% | 13.24% |
|  |  | v5.2.0 | 6.03% | 13.33% | 8.33% | 9.23% | 10.64% | 0.00% | 2.69% | 0.00% | 0.90% |
| 2,607 | psi | v5.1.0 | 28.83% | 76.67% | 75.00% | 60.17% | 60.52% | 0.00% | 16.15% | 14.48% | 10.21% |
|  |  | v5.2.0 | 1.30% | 5.13% | 12.00% | 6.14% | 3.02% | 0.00% | 2.11% | 8.98% | 3.70% |
| 3,088 | ino | v5.1.0 | 2.09% | 22.22% | 23.81% | 16.04% | 5.32% | 0.00% | 16.90% | 18.49% | 11.80% |
|  |  | v5.2.0 | 0.85% | 0.00% | 13.33% | 4.73% | 1.43% | 0.00% | 0.00% | 11.90% | 3.97% |
| 3,647 | ino | v5.1.0 | 0.63% | 17.78% | 23.26% | 13.89% | 3.90% | 0.00% | 13.88% | 19.36% | 11.08% |
|  |  | v5.2.0 | 0.22% | 0.00% | 0.00% | 0.07% | 0.94% | 0.00% | 0.00% | 0.00% | 0.00% |
| 4,305 | m<sup>5</sup>C | v5.1.0 | 13.80% | 17.65% | 4.44% | 11.97% | 0.84% | 12.96% | 16.81% | 3.60% | 11.12% |
|  |  | v5.2.0 | 3.08% | 0.00% | 2.13% | 1.73% | 1.37% | 1.71% | 0.00% | 0.76% | 1.23% |
| 5,253 | psi | v5.1.0 | 15.95% | 34.04% | 27.97% | 25.99% | 14.71% | 1.24% | 19.33% | 13.26% | 11.28% |
|  |  | v5.2.0 | 3.66% | 12.50% | 9.14% | 8.43% | 4.35% | 0.00% | 8.15% | 4.80% | 4.32% |
| 7,501 | m<sup>5</sup>C | v5.1.0 | 71.72% | 12.50% | 14.83% | 33.02% | 32.77% | 38.95% | 0.00% | 0.00% | 12.98% |
|  |  | v5.2.0 | 42.63% | 8.47% | 7.62% | 19.57% | 17.92% | 24.71% | 0.00% | 0.00% | 8.24% |
| 7,966 | ino | v5.1.0 | 7.97% | 37.60% | 37.12% | 27.57% | 21.60% | 0.00% | 16.00% | 15.52% | 10.50% |
|  |  | v5.2.0 | 9.40% | 1.27% | 1.70% | 4.13% | 2.73% | 6.67% | 0.00% | 0.00% | 2.22% |
| 8,322 | psi | v5.1.0 | 10.13% | 9.22% | 18.71% | 12.69% | 1.85% | 8.28% | 7.37% | 16.86% | 10.84% |
|  |  | v5.2.0 | 2.78% | 0.00% | 2.89% | 1.89% | 0.41% | 2.37% | 0.00% | 2.48% | 1.62% |
| 8,514 | m<sup>5</sup>C | v5.1.0 | 69.24% | 71.35% | 67.78% | 69.46% | 54.31% | 14.93% | 17.05% | 13.48% | 15.15% |
|  |  | v5.2.0 | 15.54% | 27.55% | 18.52% | 20.54% | 22.11% | 0.00% | 5.44% | 0.00% | 1.81% |
| 9,171 | psi | v5.1.0 | 25.97% | 4.66% | 8.51% | 13.05% | 0.82% | 25.15% | 3.84% | 7.69% | 12.23% |
|  |  | v5.2.0 | 1.75% | 1.39% | 1.13% | 1.42% | 0.17% | 1.58% | 1.23% | 0.96% | 1.26% |

**Table S10.**
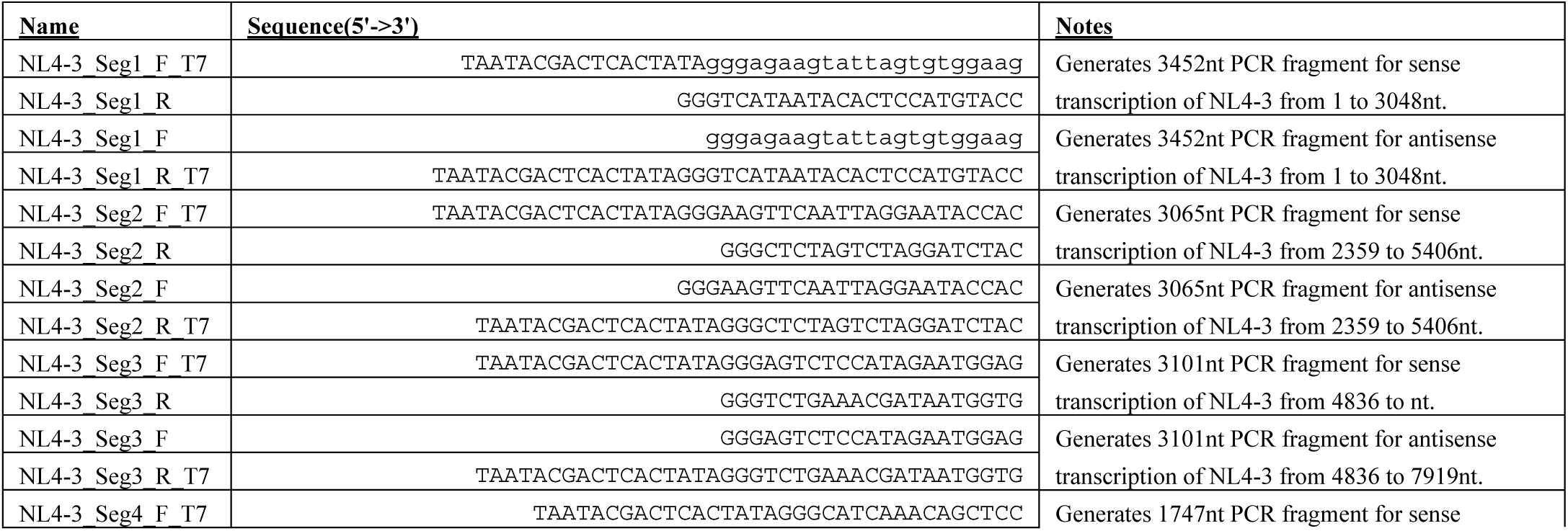
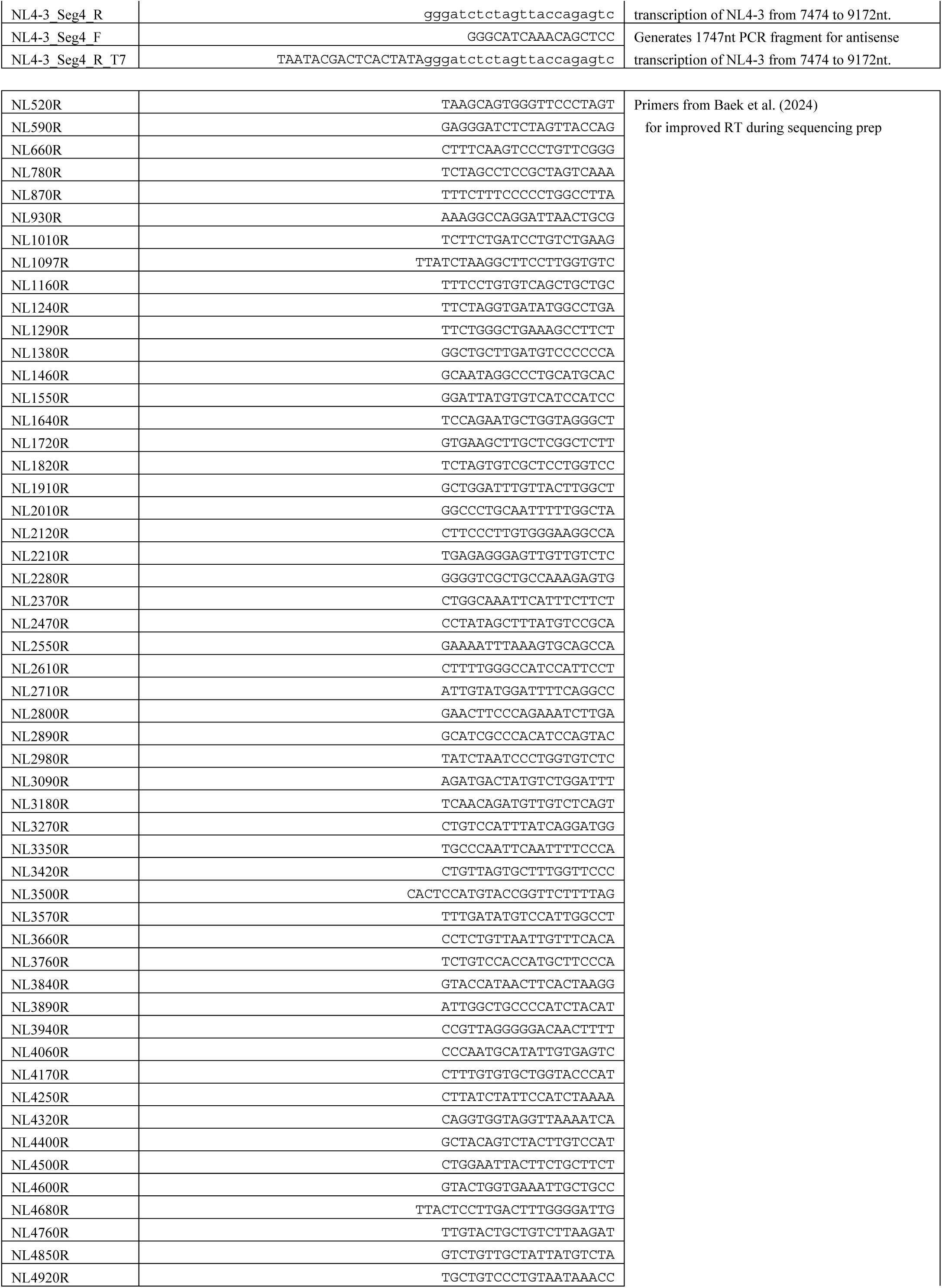
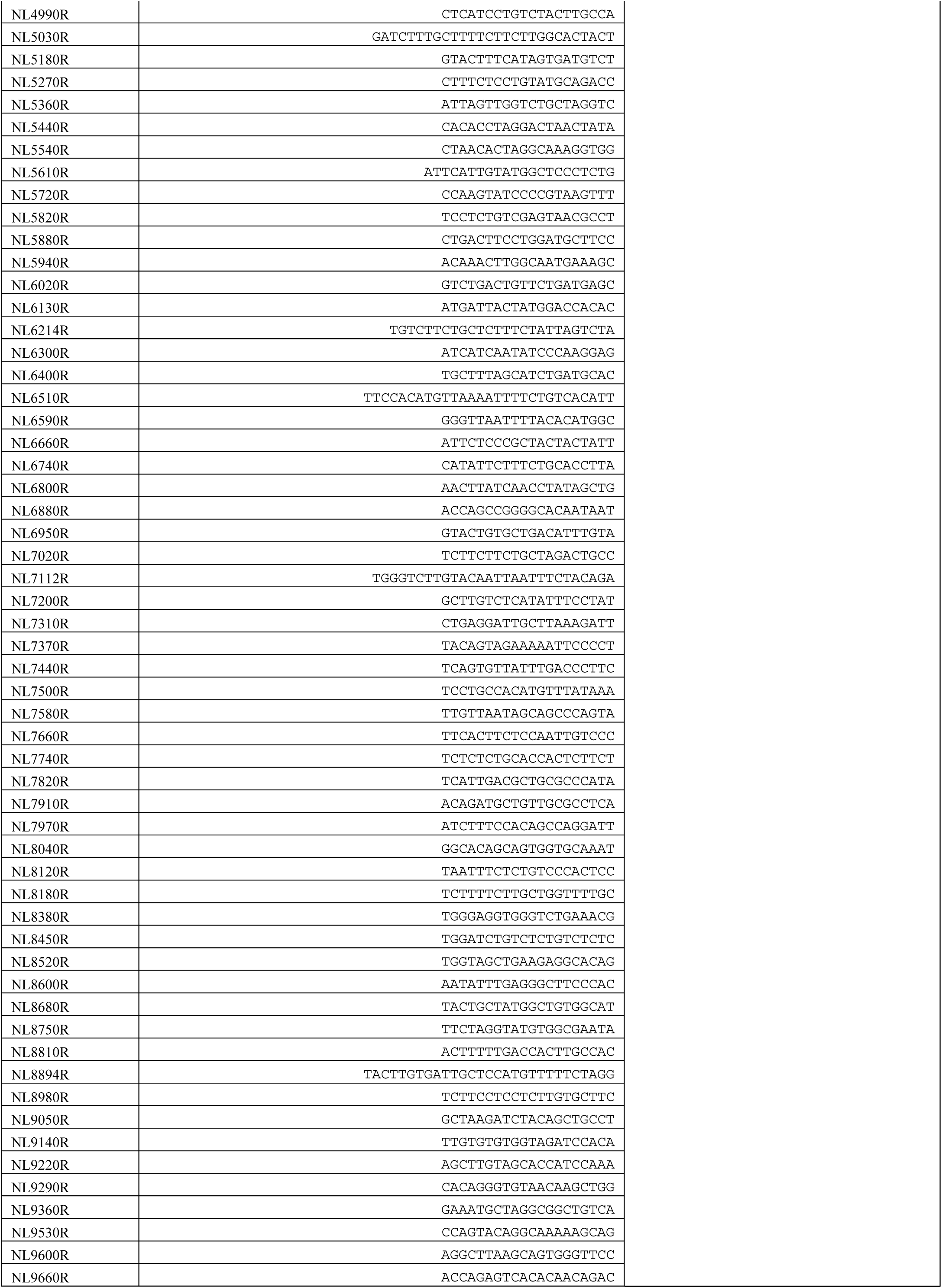
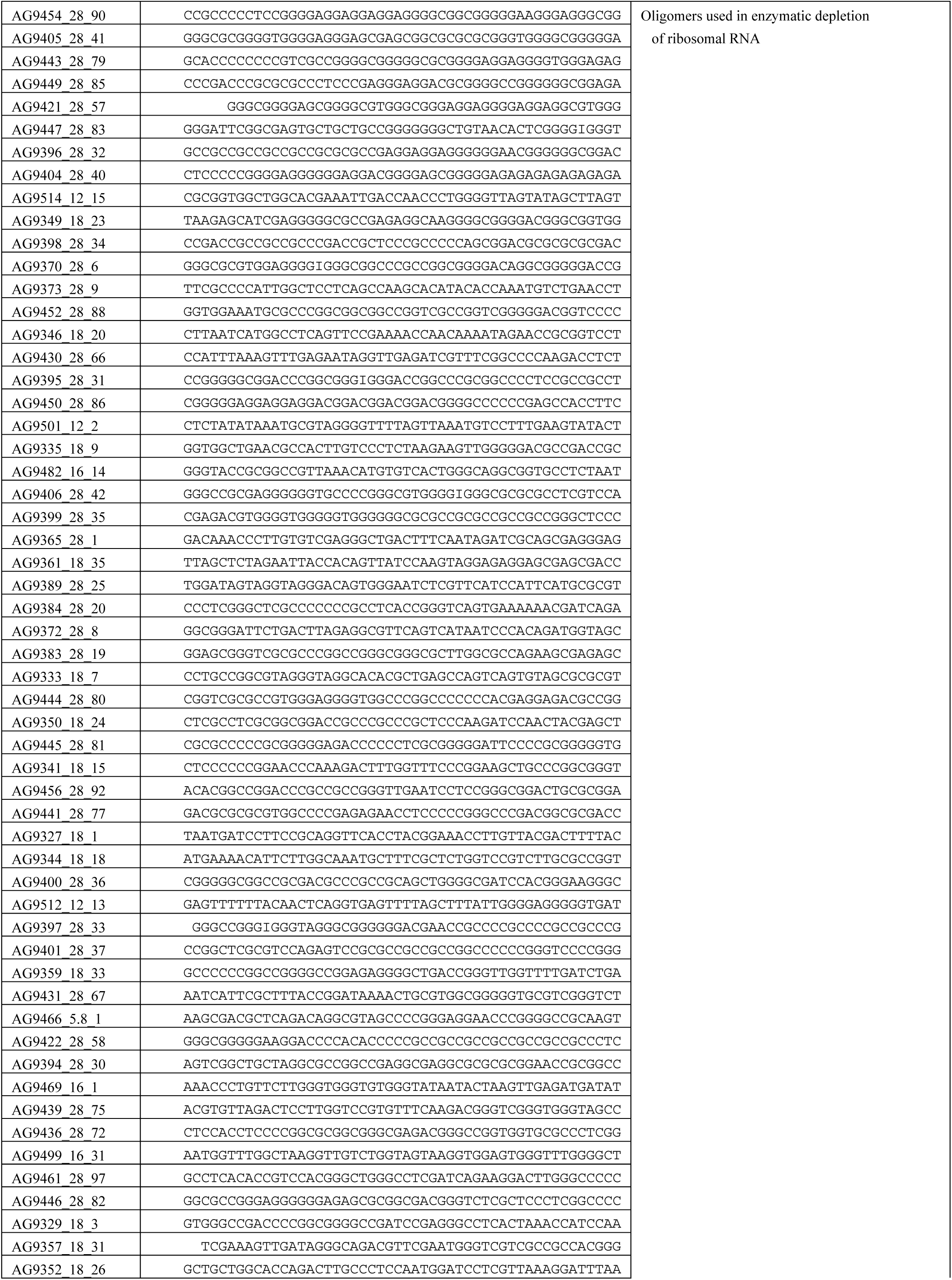
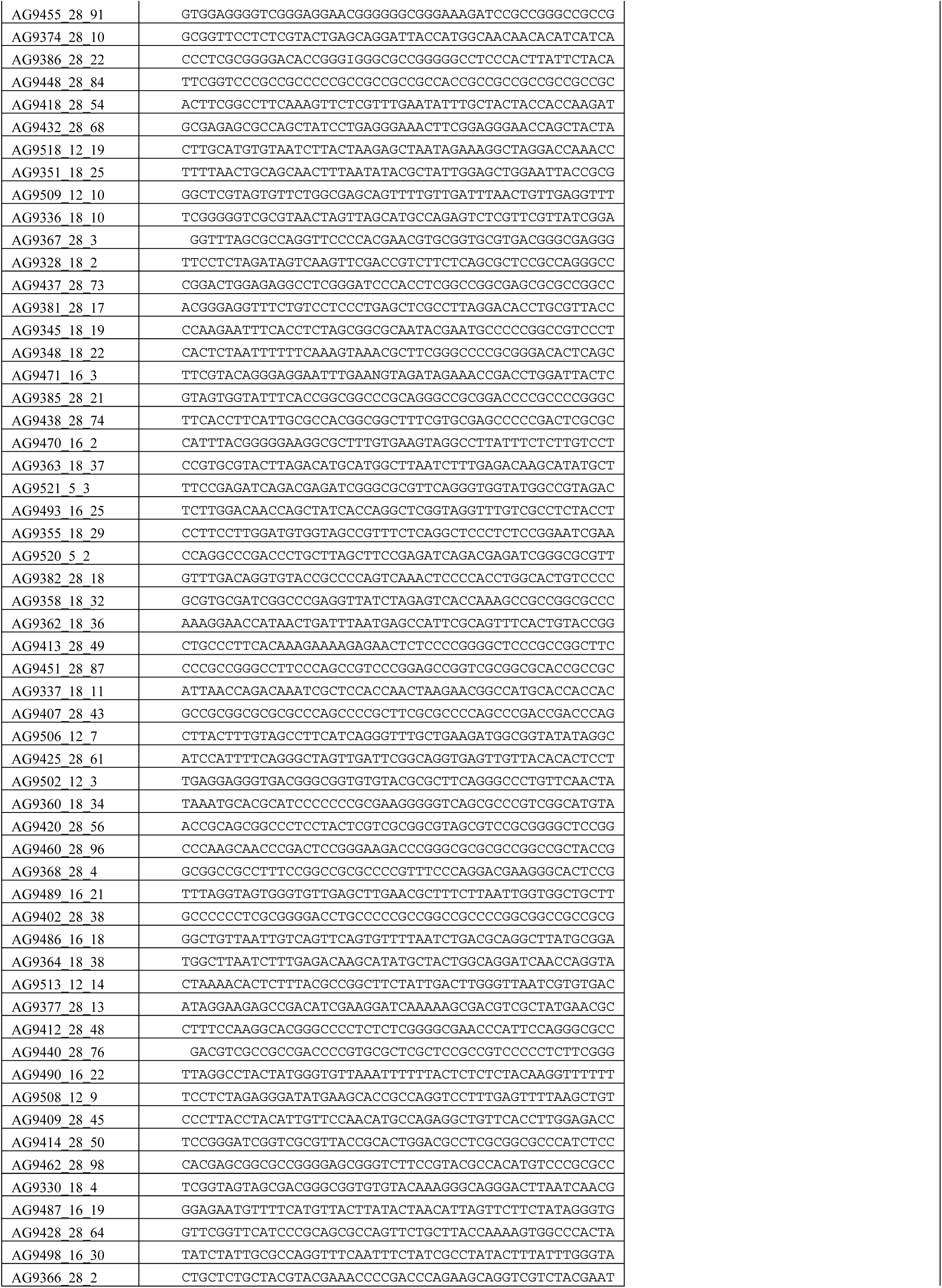
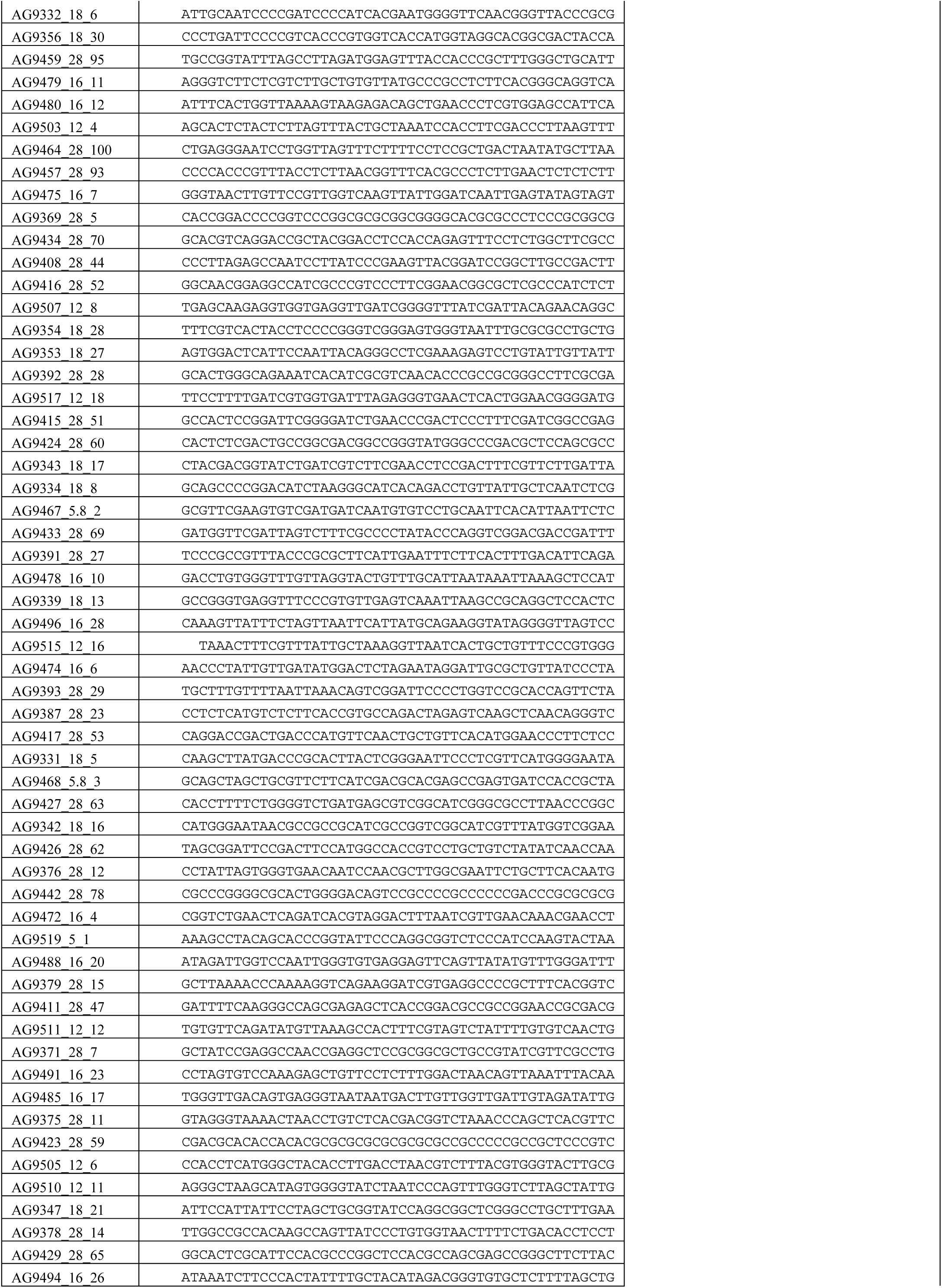
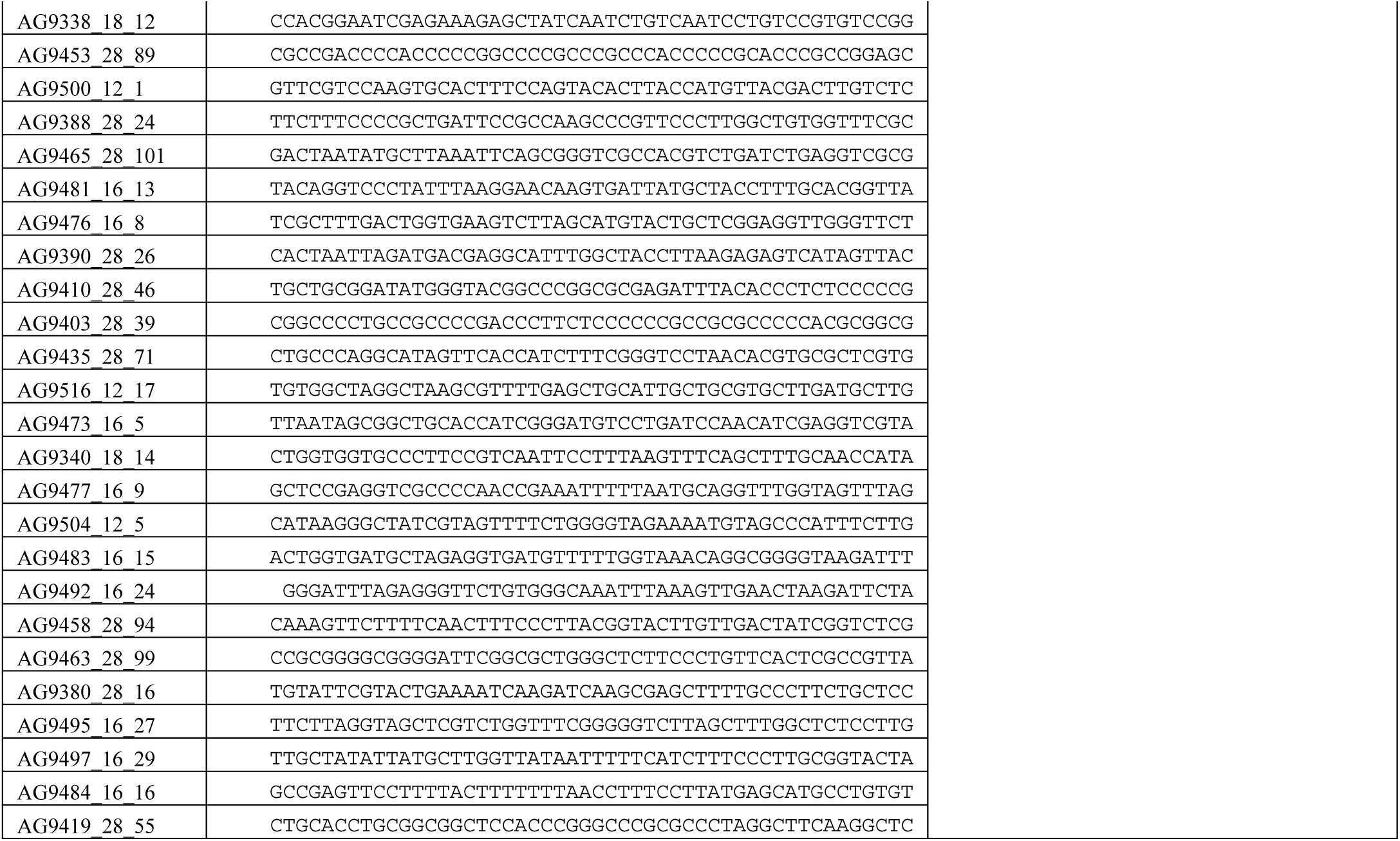
DNA oligonucleotides used in this study.

